# Dynamic Organellar Mapping in yeast reveals extensive protein localization changes during ER stress

**DOI:** 10.1101/2025.02.21.639471

**Authors:** Anna Platzek, Julia P Schessner, Klára Odehnalová, Georg H H Borner, Sebastian Schuck

**Affiliations:** Heidelberg University Biochemistry Center, Heidelberg, Germany; Department of Proteomics and Signal Transduction, Systems Biology of Membrane Trafficking Research Group, Max-Planck Institute of Biochemistry, Martinsried, Germany

**Keywords:** spatial proteomics, protein profiling, folding stress, protein relocalization

## Abstract

Sophisticated techniques are available for systematic studies of yeast cell biology. However, it remains challenging to investigate protein subcellular localization changes on a proteome-wide scale. Here, we apply Dynamic Organellar Maps (DOMs) by label-free mass spectrometry to detect localization changes of native, untagged proteins during endoplasmic reticulum (ER) stress. We find that hundreds of proteins shift between cellular compartments. For example, we show that numerous secretory pathway proteins accumulate in the ER, thus defining the extent and selectivity of ER retention of misfolded proteins. Furthermore, we identify candidate cargo proteins of the ER reflux pathway, determine constituents of reticulon clusters that segregate from the remainder of the ER and provide evidence for altered nuclear pore complex composition and nuclear import. These findings uncover protein relocalization as a major aspect of cellular reorganization during ER stress and establish DOMs as a powerful discovery tool in yeast.

## Introduction

The budding yeast, *Saccharomyces cerevisiae*, is an important model organism for basic research. Many fundamental cell biological processes, such as transcription, translation and protein targeting, have been studied in yeast through systematic approaches (DeRisi et al, 1997; Ingolia et al, 2009; Jan et al, 2014). So far, comprehensive analyses of protein localization have relied on high-throughput microscopy of large strain collections in which proteins are labeled with fluorescent tags (Huh et al, 2003; Weill et al, 2018; Meurer et al, 2018). These efforts have been complemented by automated image analysis tools (Chong et al, 2015; Kraus et al, 2017). As a result, it is now possible to compare protein localization under different conditions and thereby detect protein localization changes (Tkach et al, 2012; Breker et al, 2013; Litsios et al, 2024). Nonetheless, such systematic investigations remain complex endeavors that require substantial resources and technical expertise. Moreover, microscopy-based approaches in yeast usually involve tags, which can affect the abundance of labeled proteins, alter their localizations and disrupt their functions.

An alternative approach for investigating protein localization on a proteome-wide scale is spatial proteomics by mass spectrometry-based protein profiling (Lundberg and Borner, 2019; Borner, 2020; Christopher et al, 2021). Here, cells are lysed mechanically and fractionated by centrifugation to partially separate cell organelles. Importantly, the aim of this subcellular fractionation is not to purify individual organelles. Rather, the purpose is to generate, for each organelle, a characteristic abundance profile across fractions. Proteins in each fraction are then identified and quantified by mass spectrometry, and proteins associated with the same organelle will have similar profiles. Based on these profiles and pre-defined compartment marker proteins, machine learning techniques assign proteins to subcellular compartments to obtain an ‘organellar map’ of the cell. Comparisons of maps generated under different conditions reveal protein localization changes. To date, a single study has applied organellar mapping to yeast (Nightingale et al, 2019). This analysis investigated unperturbed cells and yielded high-confidence organelle assignments for 900 of the approximately 5400 proteins present in yeast under laboratory conditions (Ho et al, 2018). No study in yeast so far has used organellar mapping to investigate protein localization changes.

Several protein profiling methods are available, which differ in the techniques for cell lysis and fractionation, mass spectrometry and data analysis (Itzhak et al, 2016; Christoforou et al, 2016; Krahmer et al, 2018; Orre et al, 2019; Martinez-Val et al, 2021; Hein et al, 2025). One such method is the ‘Dynamic Organellar Maps’ (DOMs) approach, which has been used extensively for comparative applications (Itzhak et al, 2016; Kozik et al, 2017; Davies et al, 2018; Davies et al, 2022; Schessner et al, 2023). DOMs are based on cell fractionation by differential centrifugation and quantitative label-free mass spectrometry. The robustness and relative simplicity of these techniques ensures reproducibility and scalability, thus allowing comparisons of multiple samples. In addition, DOMs are supported by freely available online software for automated data analysis and visualization (Schessner et al, 2023). We therefore chose to adapt the DOMs approach for comparative organellar mapping in yeast.

To evaluate the utility of yeast DOMs, we investigated the cellular response to endoplasmic reticulum (ER) stress. ER stress is defined by an accumulation of misfolded newly synthesized proteins in the ER and can arise during metabolic fluctuations, adverse environmental conditions, cell differentiation and disease. ER stress activates the unfolded protein response (UPR), which enhances protein folding and the degradation of misfolded proteins (Walter and Ron, 2011). In yeast, the UPR induces hundreds of genes, which encode ER-resident protein folding and degradation machinery but also proteins that localize to post-ER compartments of the secretory pathway, mitochondria, the cytosol and the nucleus (Travers et al, 2000). Thus, ER stress alters the abundance of many proteins in various subcellular compartments. By contrast, high-throughput microscopy has identified a mere 25 proteins that showed localization changes during ER stress (Breker et al, 2013). These results raise the question whether subcellular localization changes are a minor or an underappreciated aspect of cell adaptation to ER stress.

Here, we show that DOMs detect numerous alterations of protein transport processes that are known to be affected by ER stress and additionally uncover unexpected protein localization changes in processes that have not been linked to ER stress before. These findings provide a much expanded view of proteome remodeling during ER stress and demonstrate the power of DOMs for systematic investigations in yeast.

## Results

### Establishment of the DOMs spatial proteomics approach in yeast

We first generated steady-state maps of unperturbed yeast. Cells from a liquid culture were lysed by enzymatic cell wall removal and suspension in a hypo-osmotic buffer. Organelles were released by gentle mechanical sample homogenization, unbroken cells were removed by a clearing spin, and cell lysates were fractionated through a series of centrifugation steps to obtain material that sedimented at 1, 3, 6, 12, 24 or 78 x 1,000 g, or remained in the supernatant (Figure 1A). We prepared six replicate maps, obtained in batches of two on three separate days. The total cell lysate and subcellular fractions were analyzed by quantitative label-free mass spectrometry. Protein intensity data from the total cell lysates were converted into abundance estimates in molecules per cell for 3,910 proteins (Wisniewski et al, 2014; Table S1A). The 78,000 g supernatant is called ‘cytosol fraction’, and we derived cytosolic pool estimates for each protein by dividing a protein’s intensity in the cytosol fraction by the summed intensities in all organelle and cytosol fractions (Table S1B). Of note, the cytosol fraction contained a minor proportion of soluble proteins that had leaked from the lumen of organelles and membrane proteins that presumably were part of small vesicles (Figure S1; Table S1B). Conversely, cytosolic proteins may be found in the organelle fractions if they are part of large protein complexes or have membrane-associated pools. Last, we processed the mass spectrometry data with the DOM-ABC software (Schessner et al, 2023) and identified 2,971 proteins that were reproducibly profiled across all replicates (Table S1C; Figure S2A).

**Figure 1.**
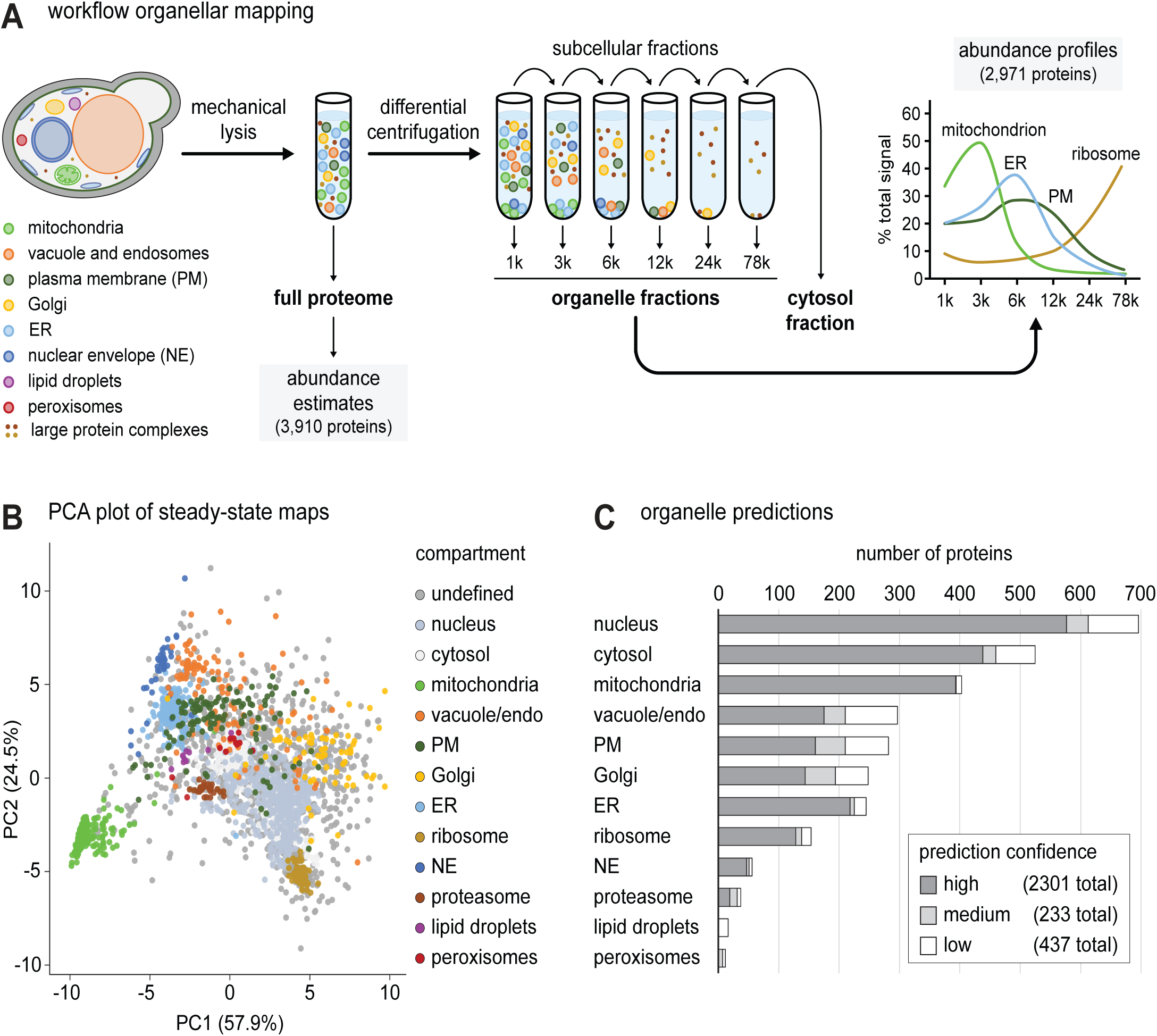
Establishment of the DOMs spatial proteomics approach in yeast. **(A)** Workflow of organellar mapping. Cells are lysed mechanically, a full proteome sample is collected and the remainder of the cell lysate is subjected to differential centrifugation to obtain subcellular fractions, including a cytosol fraction. All samples are analyzed by quantitative label-free mass spectrometry, and an abundance profile is derived for each protein. Average profiles of proteins associated with particular organelles are shown to illustrate distinct fractionation behaviors and partial separation of subcellular compartments. **(B**) Principal component analysis (PCA) plot of steady-state organellar mapping data from unperturbed yeast. PCA was performed for proteins profiled in six maps obtained in biological triplicate with two technical replicates each. Pre-defined compartment markers are colored, other proteins are classified as’undefined’. **(C)** Number of organelle predictions derived from steady-state maps with high, medium or low confidence.

To enable an assignment of the profiled proteins to cell organelles, we next defined a set of compartment marker proteins. With the help of the Saccharomyces Genome Database and relevant literature, we built a reference database of the approximately 5,400 proteins present in yeast under laboratory conditions (Kraus et al, 2017; Ho et al, 2018). Where available evidence allowed, we annotated these proteins with their predominant subcellular localization at steady state (Table S2A). Using this information, we iteratively trained the support vector machine (SVM) module of DOM-ABC (Schessner et al, 2023) to assign one of twelve possible subcellular localizations to the profiled proteins. The classifiers were nucleus, cytosol, mitochondria, vacuole (including endosomes), plasma membrane (including cell wall), Golgi, ER, ribosome, nuclear envelope, proteasome, lipid droplets and peroxisomes. Through this process, we retrieved 1,908 proteins for which the SVM assignment matched the annotation in the reference database, and we used these proteins as compartment markers (Table S2B). Visualization of the multi-dimensional map data in only two dimensions by principal component analysis (PCA) showed reproducible clustering of the compartment markers, indicating that the DOMs approach allowed robust organellar mapping in yeast (Figures 1B and S2B, C).

We then applied the trained SVMs to predict the localizations of all 2,971 profiled proteins (Figure 1C; Table S1C). This prediction showed high recall and precision across compartments (Figure S2D). For 996 proteins, the SVM prediction did not match the annotation in the reference database. This incongruence mainly arose from two sources. First, 418 proteins had classifiers in the reference database that were not available to the SVMs, such as actin-associated, ambiguous or unknown. Second, the vast majority of the remaining 578 mismatched annotations concerned proteins that had large cytosolic pools in addition to organellar or nuclear pools and thus had multiple subcellular localizations (Lundberg and Borner, 2019). In these cases, the SVM classification may not align with the majority localization in the reference database. When we disregarded proteins with classifiers unavailable to the SVMs and added ‘cytosol’ as a secondary annotation for proteins with cytosolic pool estimates >30%, the agreement between SVM predictions and the reference database increased to 89% (Table S1D). Thus, despite limitations, the SVM predictions broadly captured the complexity of subcellular protein localization and provided a useful reference grid for the interpretation of protein localization changes.

In summary, we adapted the DOMs workflow to yeast and generated protein copy number estimates, cytosolic pool estimates and subcellular localization predictions. All data can be explored through an interactive tool, which also features a ‘neighborhood predictor’ to identify proteins with similar subcellular distributions (Supplemental Database 1).

### Application of DOMs to investigate the cellular response to ER stress

To test yeast DOMs in a comparative setting, we investigated protein localization changes elicited by ER stress. To induce misfolding of newly synthesized proteins in the ER, we applied two established ER stressors: dithiothreitol (DTT), which prevents disulfide bond formation, and tunicamycin, which blocks protein N-glycosylation. We fractionated untreated, DTT-treated and tunicamycin-treated cells as above and analyzed fractions by quantitative mass spectrometry. As a prelude to organellar mapping, we determined the impact of DTT and tunicamycin on overall protein abundance. The two drugs caused distinct and reproducible proteome alterations, as indicated by PCA with global protein levels as input (Figure S3A). Volcano analysis revealed 1,457 significantly changing proteins with DTT and 1,183 with tunicamycin (Figure 2A; Table S3A, B). These numbers corresponded to about one third of the measured proteomes, indicating major proteome remodeling. We had previously defined a set of core UPR target genes (Schmidt et al, 2019). Of the corresponding proteins, most were upregulated by DTT and tunicamycin, demonstrating strong induction of ER stress (Figure 2A). These proteins included ER-resident machinery for protein folding, modification and degradation, as well as machinery for protein transport into and out of the ER. In addition, numerous post-ER secretory pathway proteins were upregulated, as were components of the proteasome (Figure S3B-E). These observations agree with earlier data and reflect that cells respond to ER stress by adjusting protein secretion and degradation. Conversely, ribosomal proteins were slightly but consistently downregulated, again in agreement with earlier data (Figure S3F; Travers et al, 2000; Schmidt et al, 2019). Hence, DTT and tunicamycin caused many protein abundance changes expected to occur during ER stress.

**Figure 2.**
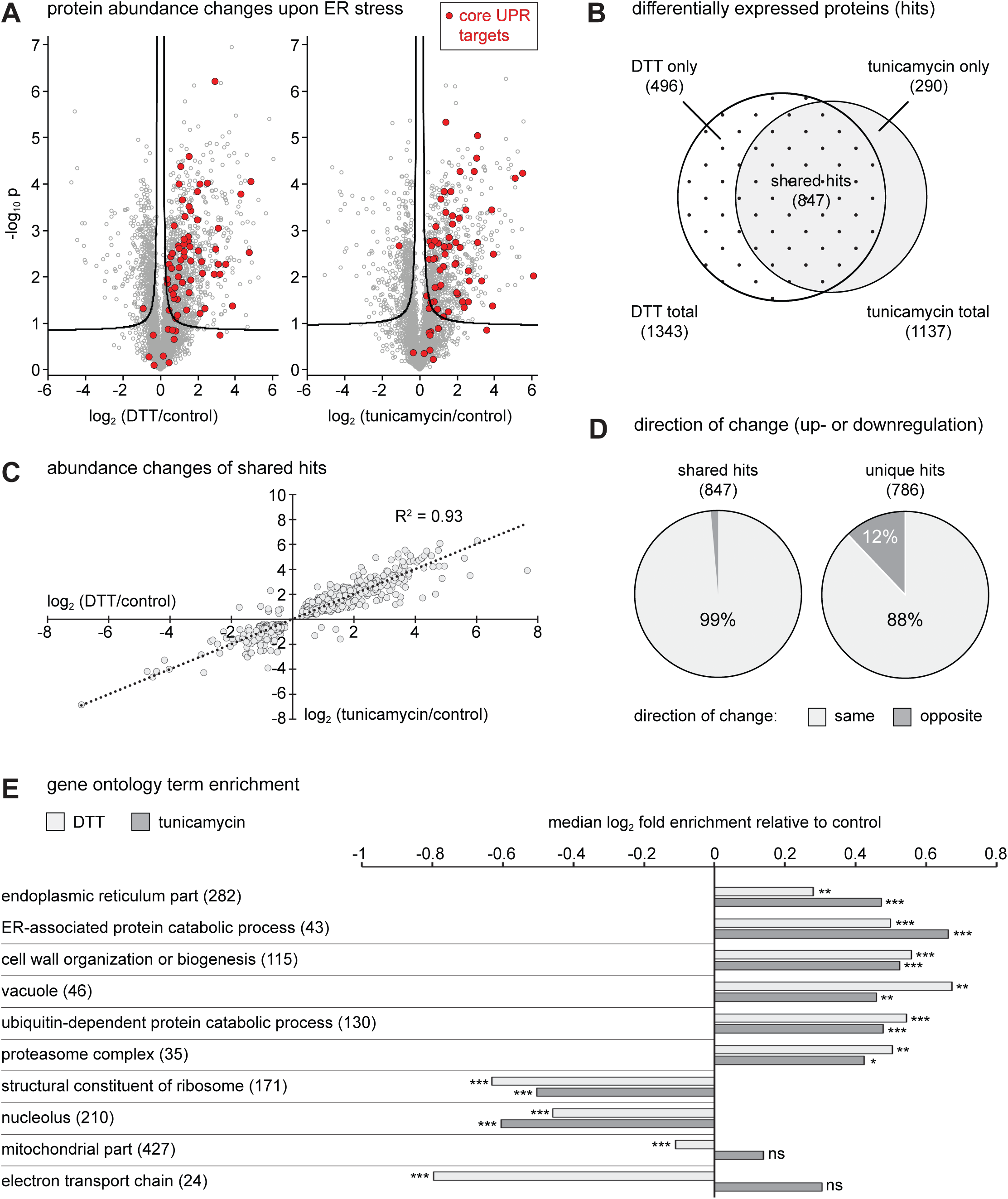
Protein abundance changes upon ER stress. **(A)** Volcano plots of full proteomes showing log_2_ fold abundance changes upon DTT or tunicamycin treatment. P-values for the significance of changes were calculated with a two-tailed t-test (n = 3). Volcano lines indicate 5% false discovery rate cut-offs based on data permutation. Core UPR targets encoded by established UPR-regulated genes are highlighted in red. **(B)** Venn diagram of differentially expressed proteins upon DTT treatment, tunicamycin treatment, or both (hits). **(C)** Scatter plot of shared hits. Plotted are the log_2_ fold abundance changes upon DTT or tunicamycin treatment. **(D)** Pie charts showing the direction of change of shared hits and proteins that qualified as hits in only one stress condition (unique hits). **(E)** Gene ontology term enrichment analysis of DTT-and tunicamycin-treated samples versus control. Selected terms show shared up-or downregulation by both treatments as well as cases of divergent regulation. The number of proteins associated with each term is given in brackets. ns, not significant; *, p<0.05; **, p<0.01; ***, p<0.001.

DTT and tunicamycin cause ER stress by different mechanisms and may have additional effects unrelated to ER stress. Of the 1,633 proteins differentially expressed in untreated and treated cells, 847 were shared between treatments, with highly correlated changes (Figure 2B, C). 99% of the 847 shared hits showed changes in the same direction (i.e. upregulated or downregulated). Remarkably, the same was true for 88% of the unique 786 hits, i.e. proteins with abundance changes that were statistically significant in only one condition (Figure 2D; Table S3C, D). Hence, most differences between the effects of DTT and tunicamycin were quantitative rather than qualitative. Still, DTT caused a specific downregulation of mitochondrial electron transport chain proteins, while tunicamycin induced a small overall upregulation of mitochondrial proteins (Figure S4A, B). Thus, mitochondria appear to be differentially affected by the two treatments. Similarly, some integral plasma membrane transporters were upregulated by DTT but downregulated by tunicamycin (Figure S4C). A gene ontology term analysis confirmed these trends (Figure 2E; Table S3E, F). In sum, the global proteome data highlighted broad similarities and specific differences in the protein abundance changes caused by DTT and tunicamycin.

Next, we analyzed protein localization changes during stress. Importantly, the abundance profiles that form the basis for assessing protein localization are normalized (see methods). As a result, localization changes can be analyzed independently of abundance changes. Organellar maps of control, DTT-treated and tunicamycin-treated cells had comparable depths of around 3,000 proteins (Figure S5A, B). PCA showed that gross map topology was similar, with two prominent exceptions (Figure 3A). First, DTT caused a striking shift of the ER cluster, which was only weakly recapitulated with tunicamycin (light blue cluster in Figure 3A, see Figure S5D for a simplified PCA plot). Second, the nuclear envelope cluster largely followed the ER shift with DTT, consistent with the physical continuity of ER and nuclear envelope (dark blue cluster in Figures 3A and S5D). Although the protein composition of the ER is similarly altered by DTT and tunicamycin (Figure 2; Reinhard et al, 2024), DTT triggers a more extensive remodeling of ER morphology (Schuck et al, 2009). This stronger impact of DTT on ER structure may result in altered vesiculation of the ER membrane upon cell lysis, a different behavior during subcellular fractionation and, hence, the shift difference revealed by PCA.

**Figure 3.**
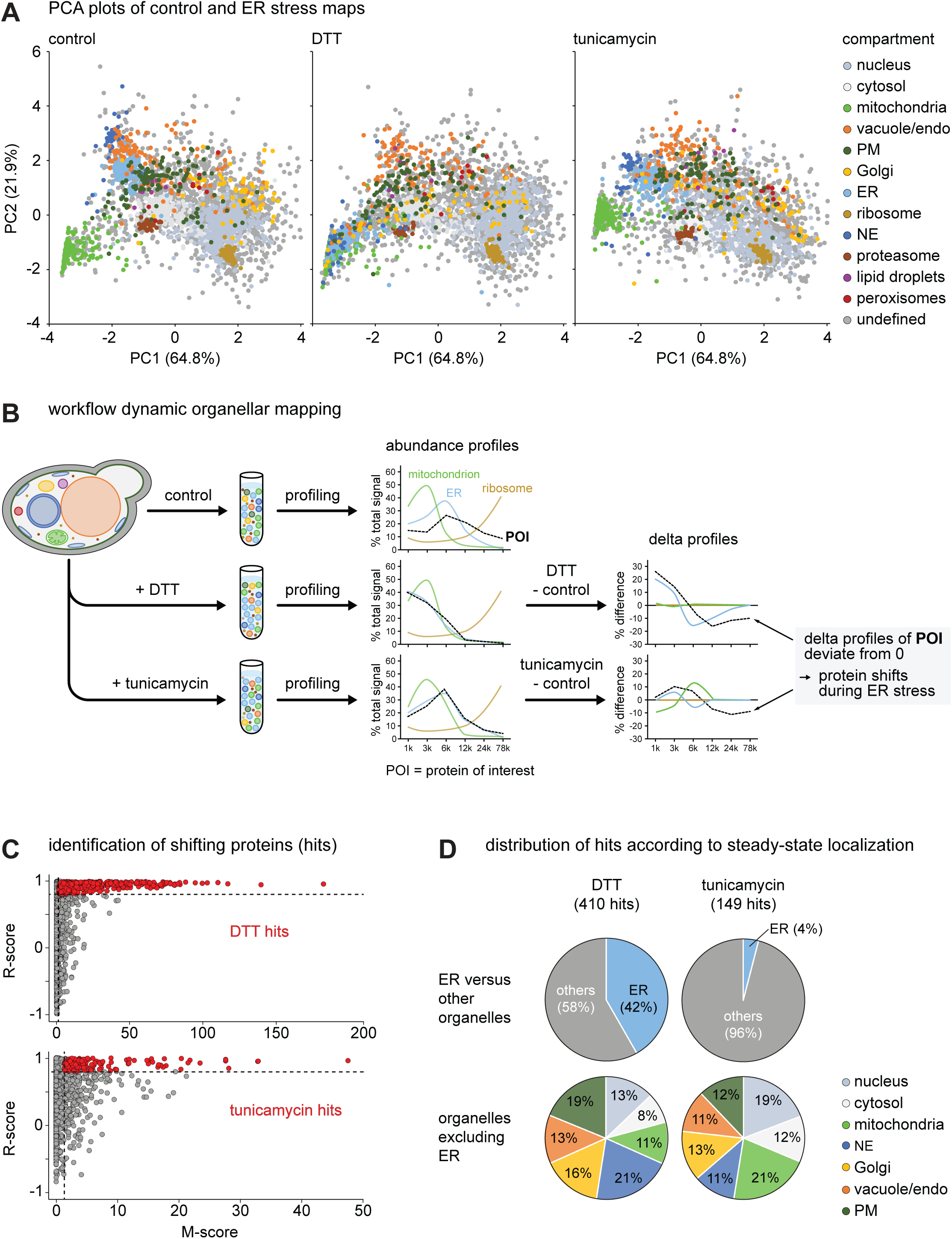
Protein localization changes upon ER stress. **(A)** Principal component analysis (PCA) plot of control and ER stress maps. Pre-defined compartment markers are colored, all other proteins are classified as’undefined’. Plots are based on the averaged profiles of three biological replicates. The most pronounced change is the shift of the ER and NE clusters (light and dark blue), which obscure the mitochondria cluster (green) in the DTT map. **(B)** Workflow of dynamic organellar mapping, illustrating the derivation of delta profiles from the comparison of abundance profiles obtained under different conditions. **(C)** Identification of shifting proteins (hits) upon DTT or tunicamycin treatment by means of movement (M) and reproducibility (R) scores. Proteins highlighted in red are candidate hits with M-scores >1.3 and R-scores >0.8, with an estimated false discovery rate of <5%. **(D)** Pie charts showing organelle distributions of DTT and tunicamycin hits according to their steady-state localization, either as ER versus other organelles (top) or among organelles excluding the ER (bottom).

We then compared control and treatment maps with the established ‘MR’ analysis to identify proteins with altered localizations (Schessner et al, 2023). This analysis is based on a comparison of a protein’s abundance profiles under control and treatment conditions (Figure 3B). Furthermore, it takes into account both the magnitude of a movement (M-score) and the reproducibility of the shift direction (R-score). At a false discovery rate of <5%, we identified 410 candidate relocalizing proteins (hits) with DTT and 149 with tunicamycin (Figure 3C; Table S4A, B). Cross-referencing with the whole proteome data revealed that the majority of these hits did not change in abundance (Figure S5C). Hence, a sizeable fraction of ER stress-responsive proteins are regulated spatially rather than by an abundance change. Analysis at the organelle level showed that DTT hits included most of the ER proteins present in the maps (Figure 3D; 171 out of 183 mapped ER proteins). This observation was consistent with the finding that the entire organelle had an altered behavior in the subcellular fractionation after DTT treatment. After excluding ER proteins, DTT and tunicamycin hits had broadly similar distributions across the remaining organelles (Figure 3D).

These data further confirmed that DTT and tunicamycin had similar effects on proteome organization. Nevertheless, DTT affected many more proteins significantly, and with higher M-scores (Figure 3C). In addition, the pronounced shifts of the ER and nuclear envelope clusters with DTT were valuable diagnostic markers for analyses of individual proteins that moved away from or towards these clusters (see below). We therefore focused our subsequent investigations on the effects of DTT.

To facilitate in-depth analyses of individual protein localization changes, we compiled all abundance and mapping data into the interactive ER Stress Maps Analysis Tool (ESMAT, Supplemental Database 2). For a given query protein, this tool provides abundance changes, cytosolic pool changes, localization predictions, shift analyses, configurable PCA and profile plots, and protein neighborhood analyses. The cytosolic pool estimates can be used to identify proteins that shift between the organelle fractions and the cytosol fraction (Table S4C). With these tools, we annotated the 410 DTT hits and categorized them into seven groups: ER proteins shifting with the whole organelle (169 proteins), proteins shifting away from the ER (12), proteins shifting towards the ER (89), nuclear envelope proteins shifting with the whole organelle (22), proteins shifting away from the nuclear envelope (17), proteins shifting away from mitochondria (7), and other localization changes (94) (Figure 4A and Table S5). Subtraction of the proteins that shifted with the whole ER or nuclear envelope left 219 proteins that showed altered distributions between compartments. For 113 of these, we were able to predict both the predominant steady-state localization and a different destination compartment, and hence the direction of a shift between two compartments (Figure 4B, C).

**Figure 4.**
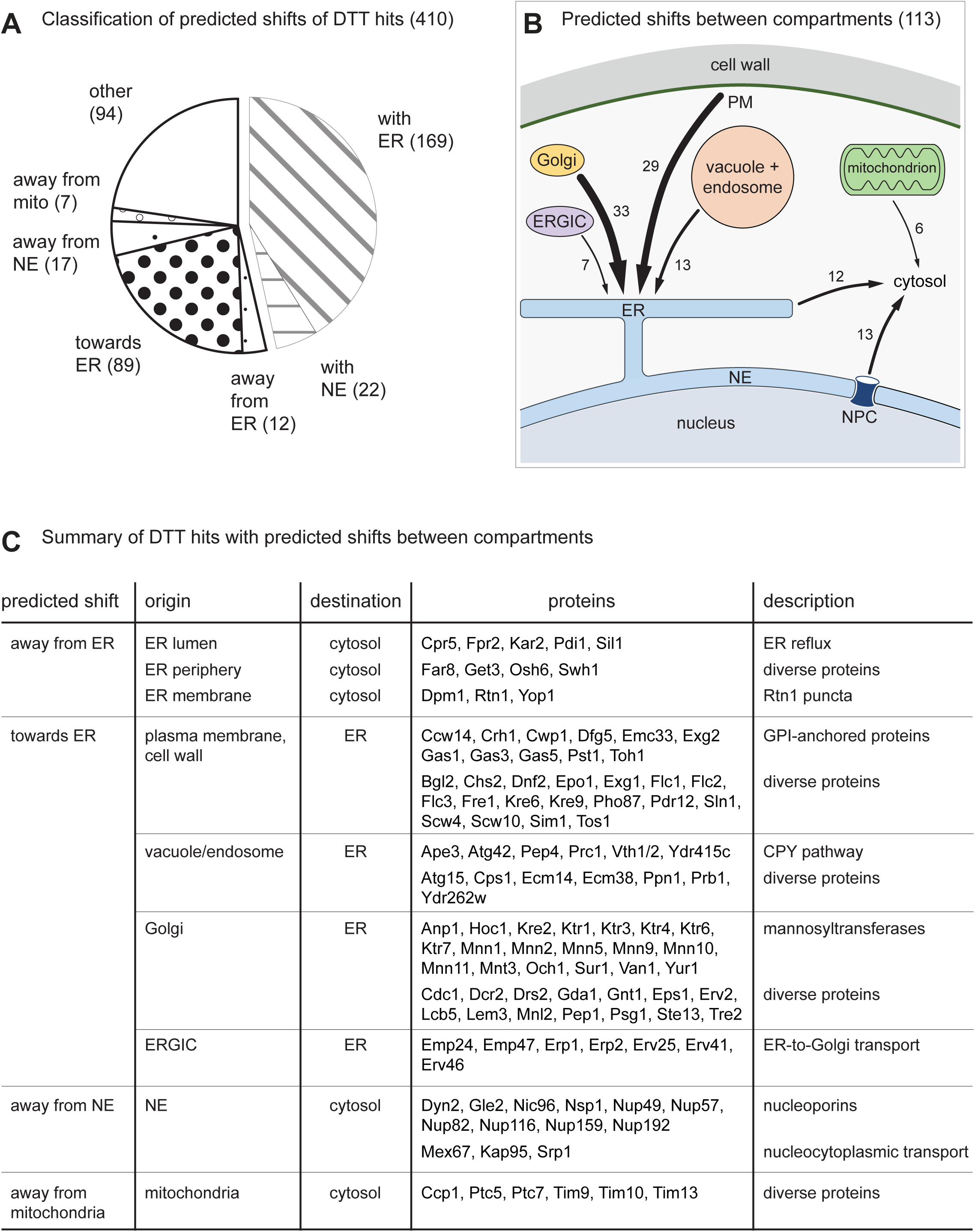
Predicted protein localization changes upon DTT-induced ER stress. **(A)** Pie chart showing classification of predicted shifts of DTT hits, including shifts with the whole ER or nuclear envelope (NE). The number of proteins per category is given in brackets. mito, mitochondria. **(B)** Illustration of predicted shifts between compartments for 113 DTT hits. ERGIC, ER-Golgi intermediate compartment; PM, plasma membrane; NPC, nuclear pore complex. **(C)** Summary of DTT hits with predicted shifts between compartments.

### Experimental evaluation of candidate relocalizing proteins

We next tested the validity of the above analysis by microscopy. For this purpose, we fused candidate relocalizing proteins to fluorescent proteins through chromosomal gene tagging. Depending on the protein analyzed, we positioned the tag such that determinants important for correct localization remained intact, such as signal sequences for ER import, ER retrieval sequences, transmembrane domains of tail-anchored proteins, and glycosylphosphatidylinositol (GPI) anchor attachment sequences. For some candidates, the tag had to be placed into the lumen of an organelle, either because the tagged protein was entirely lumenal or because tagging of a transmembrane protein at its cytosolic terminus would have caused mislocalization. In these cases, we tagged proteins with superfolder GFP (sfGFP), which matures more effectively in the oxidizing interior of the ER than other fluorescent proteins, including the original GFP (Aronson et al, 2011).

### Proteins moving away from the ER

The 12 proteins moving away from the ER significantly shifted towards the cytosol upon both DTT and tunicamycin treatment (Figure 5A and Table S5). Five of these were soluble lumenal ER proteins, namely Cpr5, Sil1, Fpr2, Pdi1 and Kar2. Soluble proteins that have been targeted correctly to the ER lumen can subsequently relocalize to the cytosol during stress. This reverse translocation, called ER reflux, is functionally enigmatic and has been analyzed mainly with artificial reporters. However, it has been demonstrated also for the endogenous proteins Cpr5 and Pdi1 (Igbaria et al, 2019; Lajoie et al, 2020). Of the over 300 ER proteins present in yeast, only 19 are lumenal (Table S2C), and the proteomic data permitted an analysis of all of them. Besides the five proteins mentioned above, only Lhs1, Eug1, Scj1 and Mpd1 showed weak evidence of ER reflux, and the remaining ten lumenal proteins had no detectable cytosolic pools (Figure 5B). This selectivity indicated that the ER membrane remains intact during cell fractionation and suggested that only certain ER proteins are reflux cargos. Fluorescence microscopy confirmed that sfGFP-labeled Sil1 and Pdi1 redistributed to the cytosol during DTT treatment, whereas Ero1 did not (Figures 5C and S6A). Overall Ero1 levels rose 8-fold during stress and thus more steeply than the levels of, for example, Cpr5, Fpr2, Pdi1 and Kar2 (Table S3A). Hence, ER import of newly synthesized Ero1 remained intact during stress, arguing against the possibility that the cytosol shifts of Cpr5, Sil1, Fpr2, Pdi1 and Kar2 were due to defects in protein targeting to the ER. Overall, DOMs indicate that ER reflux is a selective transport process and provide an extended, and possibly complete, list of cargo proteins.

**Figure 5.**
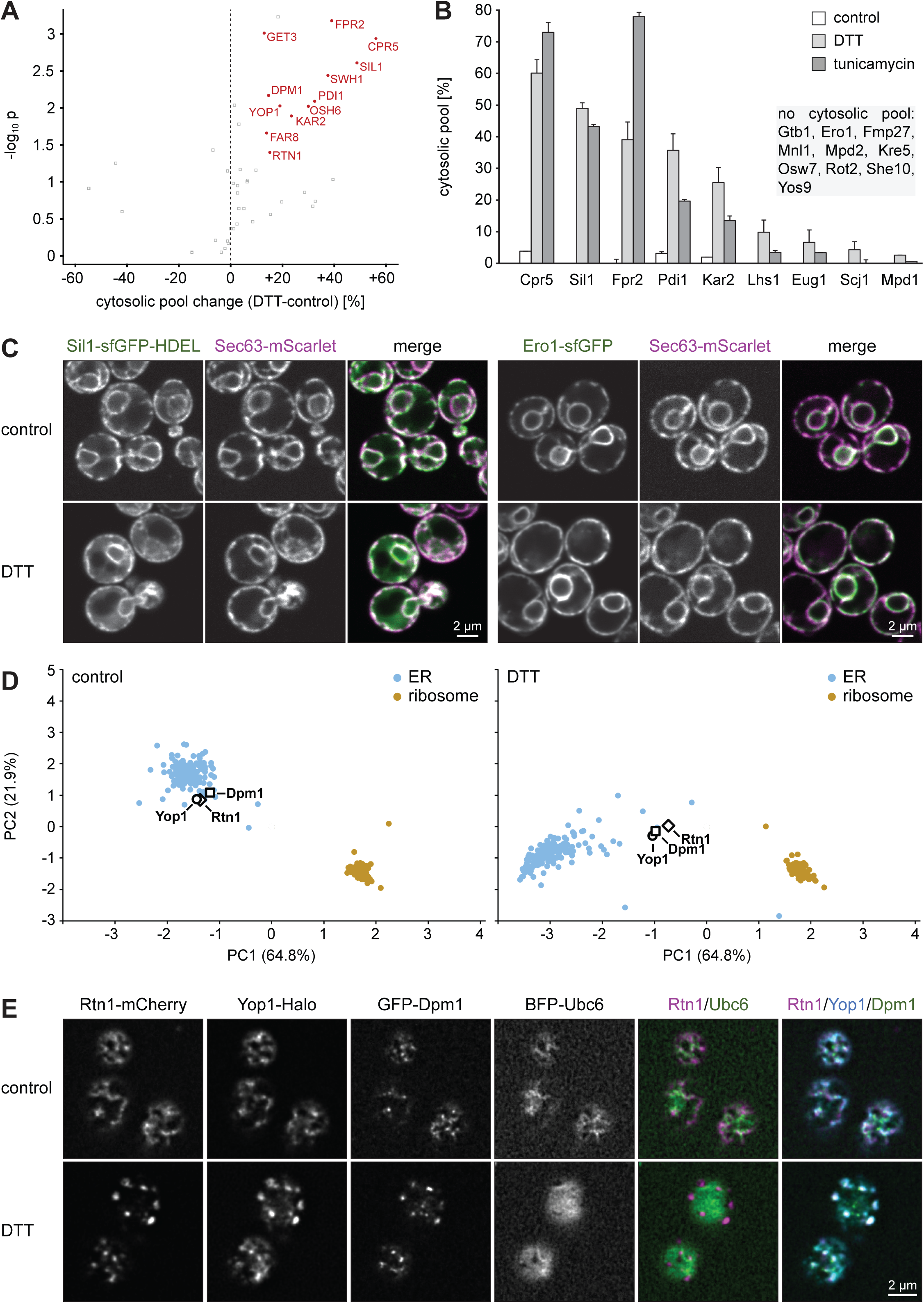
ER reflux and reticulon clustering upon ER stress. **(A)** Volcano plot of the % change of the cytosolic pool of ER proteins upon DTT treatment. P-values for the significance of change were calculated with a two-tailed paired t-test (n = 3). Proteins with a consistent increase of their cytosolic pool of >10% with both DTT and tunicamycin treatment are highlighted in red. **(B)** Cytosolic pools of lumenal ER proteins in control, DTT-treated and tunicamycin-treated cells. Bars represent the mean of three biological replicates, error bars show standard deviations. **(C)** Confocal fluorescence images of mid sections of control and DTT-treated cells expressing the ER marker Sec63-mScarlet along with the lumenal ER proteins Sil1-sfGFP-HDEL or Ero1-sfGFP. Sil1 becomes partially cytosolic during ER stress, Ero1 remains ER-localized. **(D)** PCA plot of organellar maps of control and DTT-treated cells. Rtn1, Yop1 and Dpm1 shift away from the ER cluster upon DTT treatment. The ribosome cluster does not shift and is shown for reference. **(E)** Deconvolved confocal fluorescence images of cortical sections of control and DTT-treated cells expressing the ER marker BFP-Ubc6 along with Rtn1-mCherry, Yop1-Halo and GFP-Dpm1. Rtn1, Yop1 and Dpm1 co-cluster upon DTT treatment.

Besides the lumenal ER proteins discussed above, four peripheral ER membrane proteins (Get3, Swh1, Osh6, Far8) and three integral ER membrane proteins (Rtn1, Yop1, Dpm1) shifted towards the cytosol (Figure 5A). A cytosolic shift of integral membrane proteins was unexpected. PCA showed that Rtn1, Yop1 and Dpm1 segregated from the ER upon stress and clustered tightly (Figure 5D). We found previously that Rtn1 forms punctate structures upon ER stress, and similar structures have been observed for a Yop1 homolog in fission yeast (Papagiannidis et al, 2021; Wang et al, 2023). Rtn1 and Yop1 are reticulon homology domain proteins important for ER morphogenesis (Voeltz et al, 2006). They interact with each other and with the tail-anchored protein Dpm1 (Mast et al, 2016). Four-color imaging showed that Rtn1, Yop1 and Dpm1 indeed co-localized in DTT-induced puncta, which segregated from the general ER marker Ubc6 (Figure 5E). The nature of the underlying structures remains to be clarified. In particular, it will be interesting to determine whether Rtn1, Yop1 and Dpm1 are part of a stress-induced ER subdomain or detach from the ER during stress. Either scenario could explain their shift to the cytosol fraction, which likely contains also small membrane vesicles that form naturally or arise during cell fractionation. In any case, DOMs identified new components of a largely uncharacterized ER subdomain or ER-derived structure.

### Proteins redistributing towards the ER

It has long been appreciated that newly synthesized proteins that fail to fold properly after import into the ER are retained there, thus ensuring that only functional proteins are supplied to post-ER compartments (Rose and Doms, 1988; Hurtley and Helenius, 1989). The stressors DTT and tunicamycin cause pervasive protein misfolding and are expected to lead to ER retention of many proteins. The DOMs approach made it possible to test this expectation on a proteome-wide scale. To identify proteins that redistributed to the ER during stress, we searched for candidates whose abundance profiles across subcellular fractions became more similar to the average profile of ER marker proteins. The large baseline shift of the ER cluster upon DTT treatment strongly enhanced the predictive power of this criterion. In addition, we required that candidates were also flagged as hits in the MR analysis and had a known or predicted post-ER secretory pathway localization. This analysis yielded 89 hits (Table S5). In untreated cells, 7 of these localized to the ER-Golgi intermediate compartment (ERGIC), 33 to the Golgi, 29 to the plasma membrane or cell wall, 13 to the vacuole or endosomes, and 7 had an unclear post-ER localization in the secretory pathway. Of these 89 hits, 86 had predicted transmembrane domains or ER-targeting signal peptides and likely are integral membrane or lumenal proteins. The mapped and MR-analyzed proteins in DTT-treated cells included 243 integral membrane or lumenal proteins that localize to post-ER secretory pathway compartments (Table S4E). These proteins must therefore pass through the ER. The 86 transmembrane or lumenal proteins shifting to the ER are a minority among these 243 proteins. This fact implies that there was no general ER export block during stress but that certain proteins traversing the ER were particularly liable to misfolding and retention.

The 13 vacuole or endosome proteins shifting towards the ER included all five soluble proteins that were present in the maps and are targeted to the vacuole lumen via the CPY pathway (Prc1/Cpy, Pep4, Atg42, Ape3, Ydr415c; Eising et al, 2022). In addition, they included two membrane proteins that act as sorting receptors in the CPY pathway (Vth1 and Vth2; Cooper and Stevens, 1996). Pep1/Vps10, the third sorting receptor of the pathway, cycles between the Golgi and endosomes (Marcusson et al, 1994), and it also shifted towards the ER upon stress (Figure 6A; Table S5). By contrast, transport of vacuole membrane proteins along the ALP pathway appeared to remain intact because none of its prototypical cargos changed their predicted localization (Pho8, Nyv1, Yck3, Atg27; Eising et al, 2022; Table S4A). Prc1 shifted particularly strongly as its abundance profile correlated well with the average profile of vacuole marker proteins under control conditions but aligned with the average profile of ER marker proteins under stress (Figure 6B). Microscopy confirmed DTT-induced redistribution of Prc1, Atg42 and Pep1 towards the ER (Figures 6C and S6B). Since Prc1 needs to reach the vacuole to become active, this localization change explains why Prc1 activity declines during ER stress, even though its abundance increased three-fold (Schuck et al, 2014; Table S3A). These results indicate that ER stress impairs the targeting of vacuole proteins by causing misfolding and ER retention of cargos and sorting receptors, particularly of the CPY pathway.

**Figure 6.**
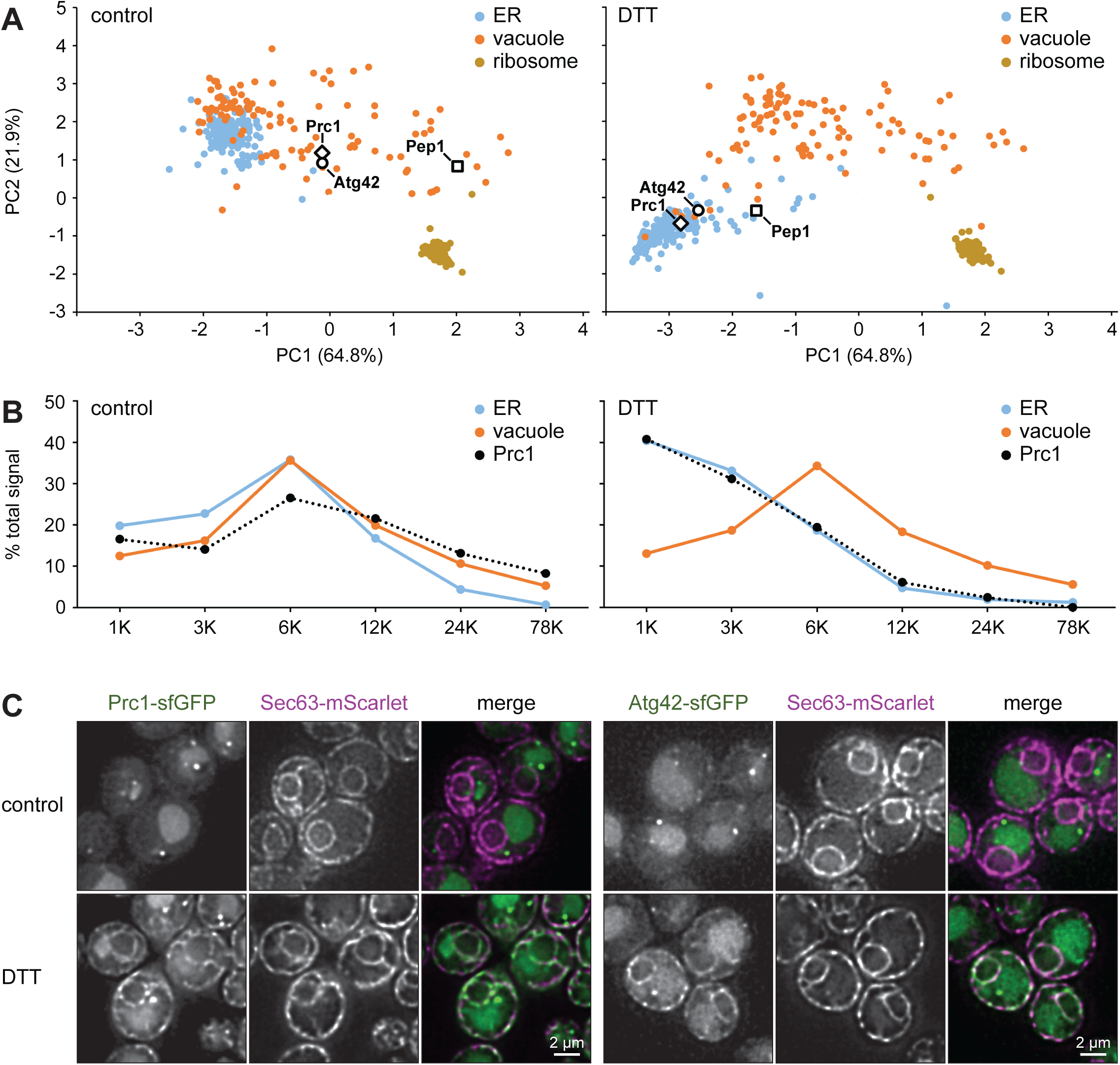
Redistribution of vacuole proteins towards the ER upon ER stress. **(A)** PCA plot of organellar maps of control and DTT-treated cells. Prc1, Atg42 and Pep1 shift towards the ER cluster upon DTT treatment. The ribosome cluster is shown for reference. **(B)** Abundance profile of Prc1 and average abundance profiles of ER and vacuole marker proteins in control and DTT-treated cells. The Prc1 profile correlates with the vacuole profile in control cells and aligns with the ER profile upon DTT treatment. **(C)** Deconvolved widefield fluorescence images of mid sections of control and DTT-treated cells expressing the ER marker Sec63-mScarlet along with the vacuole proteins Prc1-sfGFP or Atg42-sfGFP. Prc1 and Atg42 localize to endosomes (bright puncta) and the vacuole at steady state but redistribute towards the ER upon DTT treatment.

The 29 plasma membrane or cell wall proteins with stress-induced shifts towards the ER included 11 predicted or validated GPI-anchored proteins (out of 13 mapped GPI-anchored proteins; Hamada et al, 1998; Pittet and Conzelmann, 2007). GPI-anchored proteins are exposed to the extracellular space and are expected to be post-translationally modified. It is therefore plausible that their maturation requires the ER-resident protein folding and modification machinery, which becomes overwhelmed during stress. Gas1 and Gas3 were two GPI-anchored proteins that showed large shifts towards the ER cluster in PCA plots (Figure 7A). Microscopy confirmed their DTT-induced redistribution towards the ER (Figures 7B and S6C). In addition, we tested Utr2, which was one of the two GPI-anchored proteins that had not satisfied all of our criteria for proteins shifting towards the ER (Table S4A). Nevertheless, Utr2 showed a moderate shift in PCA plots (Figure 7A) and appeared in the ER upon DTT treatment (Figure 7B, note the Utr2 signal at the nuclear envelope). These results suggested that most GPI-anchored proteins were subject to ER retention during DTT-induced stress. Next, we validated ER redistribution of Golgi proteins. Of the 33 proteins suggested by the data analysis, 19 were mannosyltransferases (out of 20 Golgi-localized mannosyltransferases present in the maps; Figure 7C; Fabre et al, 2014). We confirmed the predicted DTT-induced ER relocalization for Mnn2 and Mnn5 (Figures 7D and S6D). By contrast, and as predicted by the maps, the Golgi membrane protein Aur1 did not relocalize, showing that only specific Golgi proteins accumulated in the ER (Figure 7D). It remains to be determined whether mannosyltransferases accumulate in the ER because of the misfolding of newly synthesized molecules or because they naturally cycle between the ER and the Golgi, and stress shifts the transport equilibrium towards the ER (Todorow et al, 2000).

**Figure 7.**
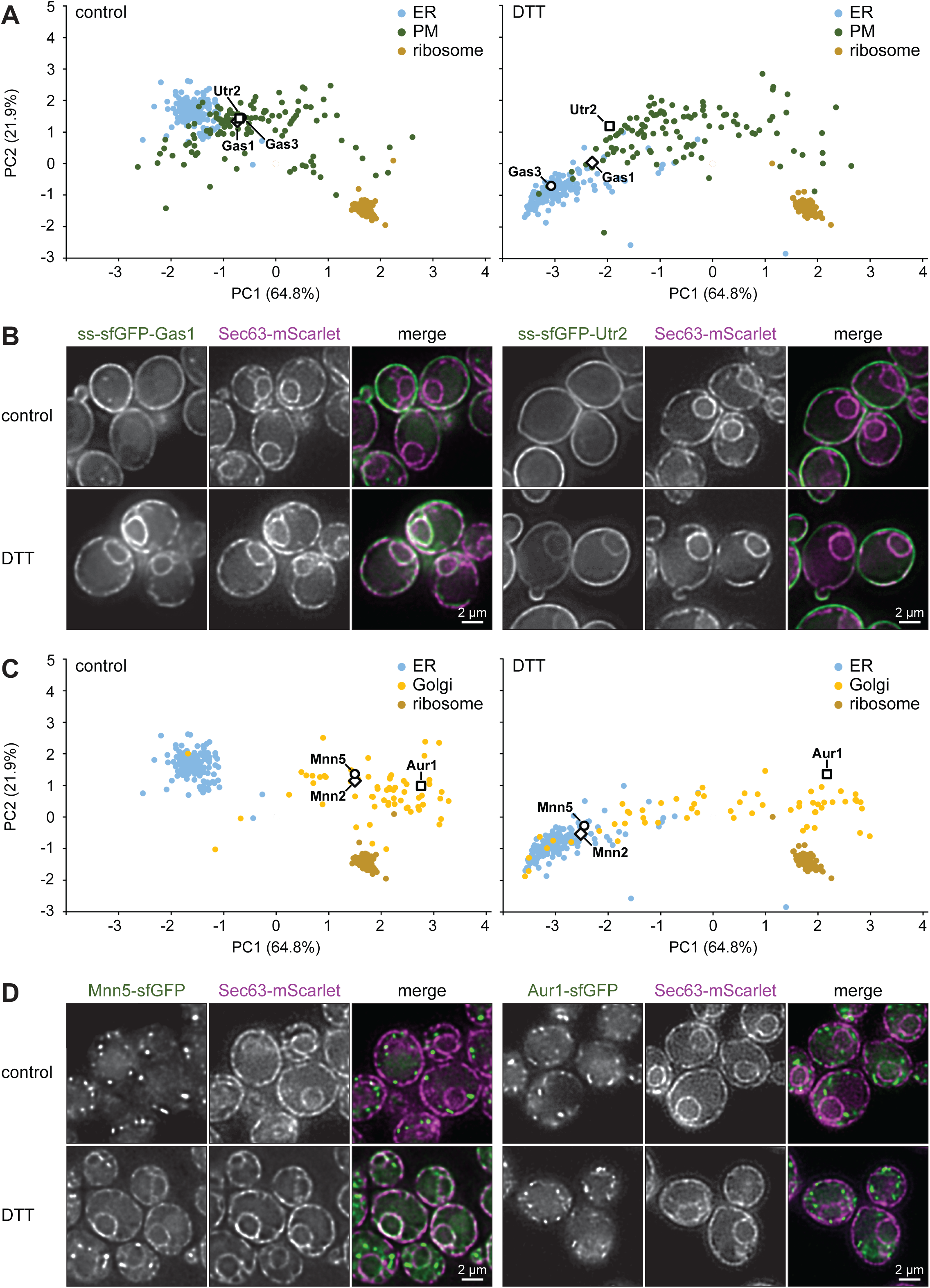
Redistribution of plasma membrane and Golgi proteins towards the ER upon ER stress. **(A)** PCA plot of organellar maps of control and DTT-treated cells. The GPI-anchored proteins Gas1, Gas3 and Utr2 shift towards the ER cluster upon DTT treatment. The ribosome cluster is shown for reference. **(B)** Deconvolved widefield fluorescence images of mid sections of control and DTT-treated cells expressing the ER marker Sec63-mScarlet along with ss-sfGFP-Gas1 or ss-sfGFP-Utr2 (ss = signal sequence for ER targeting). Gas1 and Utr2 redistribute from the plasma membrane towards the ER upon DTT treatment. **(C)** PCA plot of organellar maps of control and DTT-treated cells. The Golgi proteins Mnn2 and Mnn5 but not Aur1 shift towards the ER cluster. **(D)** Deconvolved widefield fluorescence images of mid sections of control and DTT-treated cells expressing the ER marker Sec63-mScarlet along with Mnn5-sfGFP or Aur1-sfGFP. Mnn5 but not Aur1 redistributes from the Golgi towards the ER upon DTT treatment.

Taken together, DOMs identified many secretory pathway proteins that accumulated in the ER, either due to misfolding and ER retention of newly synthesized molecules or due to trapping of pre-existing molecules that normally cycle within the early secretory pathway. These results agree with prior knowledge of ER protein quality control and reveal the extent to which stress alters protein transport. Furthermore, they indicate that specific classes of secretory proteins, namely lumenal vacuole proteins, GPI-anchored proteins and Golgi-localized mannosyltransferases, are particularly liable to mislocalization.

### Redistribution of nuclear pore complex components and importins

Finally, we turned to subcellular localization changes unrelated to the secretory pathway. The MR analysis indicated significant localization changes for many nuclear envelope proteins, including most components of the nuclear pore complex. However, a closer look at nuclear pore complex components (called nucleoporins) uncovered a dichotomy. While 15 nucleoporins shifted with the nuclear envelope cluster, 10 shifted away from the main cluster (Figure 8A, compare Nsp1, Nup57, Nup82 with Nup84, Nup133, Nup170). Microscopy revealed that these divergent behaviors corresponded to distinct subcellular distributions, as Nsp1 formed cytosolic puncta upon ER stress whereas Nup84 did not (Figure 8B). We then imaged all 31 nucleoporins. Strikingly, the 10 nucleoporins predicted to shift away from the nuclear envelope formed cytosolic puncta, whereas the 15 nucleoporins predicted to shift with the nuclear envelope did not. Of the remaining six nucleoporins that were not DTT hits in the MR analysis, another two formed cytosolic puncta (Table S6). Mapping these imaging results onto the PCA plot of DTT-treated cells revealed a clear separation of puncta-forming from non-puncta-forming nucleoporins (Figure 8C). This separation corresponded well with the known subcomplexes of the nuclear pore complex (Beck and Hurt, 2017; Akey et al, 2022; Dultz et al, 2022). Specifically, components of the cytoplasmic filaments and the channel complex as well as some linker Nups and inner ring components formed puncta. By contrast, components of the nuclear basket, membrane ring and outer ring did not form puncta (Figure 8D, puncta-forming nucleoporins are underlined). Hence, DOMs accurately identified a subset of nucleoporins that partially redistributed to the cytosol upon stress.

**Figure 8.**
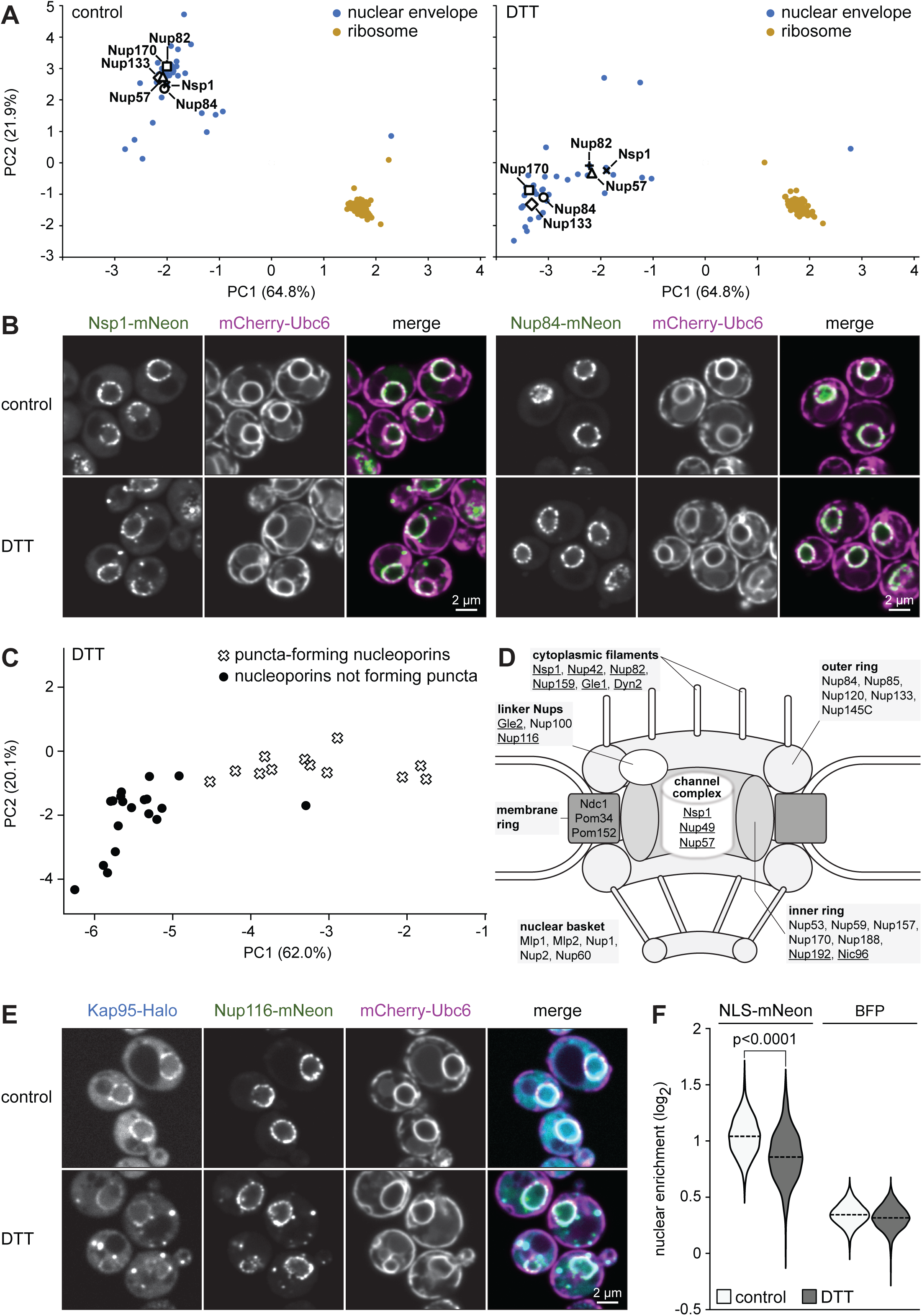
Impact of ER stress on the localization of nuclear pore complex components and importins. **(A)** PCA plot of organellar maps of control and DTT-treated cells. Nucleoporins shift with the nuclear envelope cluster (e.g. Nup84) or segregate from it (e.g. Nsp1). The ribosome cluster is shown for reference. **(B)** Confocal fluorescence images of mid sections of control and DTT-treated cells expressing the ER marker mCherry-Ubc6 along with Nsp1-mNeon or Nup84-mNeon. Nsp1 forms cytosolic puncta upon DTT treatment but Nup84 does not. **(C)** PCA plot of organellar maps of DTT-treated cells showing segregation of puncta-forming nucleoporins from nucleoporins that do not form cytosolic puncta. **(D)** Schematic architecture of the nuclear pore complex showing seven subcomplexes. Nucleoporins that form cytosolic puncta upon DTT treatment are underlined. **(E)** Confocal fluorescence images of mid sections of control and DTT-treated cells expressing the ER marker mCherry-Ubc6 along with the importin Kap95-Halo and the nucleoporin Nup116. **(F)** Nuclear import of mNeon with a nuclear localization sequence (NLS-mNeon) expressed from an inducible promoter system for 50 min. The plot shows the log_2_ fold enrichment of NLS-mNeon in nuclei versus whole cells in control and DTT-treated cells. Constitutively expressed BFP without a targeting sequence served as a control for passive accumulation in the nucleus. Dashed lines indicate sample medians. For each condition, ≥800 cells from three biological replicates were quantified. The p-value was calculated with a two-tailed Mann-Whitney U-test. Nuclear enrichment of NLS-mNeon was reduced in DTT-treated cells. Nucleo-cytoplasmic distribution of BFP was virtually unaffected by DTT, although the marginal difference between control and DTT-treated cells also reached statistical significance due to large sample sizes.

Intriguingly, the puncta-forming nucleoporins Nup116 and Gle2 and the importin Kap95 showed very similar DTT-induced shifts on PCA plots (Figure S7A). Nup116 and Gle2 are known to interact, as are Nup116 and Kap95 (Iovine et al, 1995; Bailer et al, 1998). The common shift of Nup116 and Kap95 corresponded to the formation of cytosolic puncta containing both proteins (Figure 8E). Given this cytosolic sequestration of the nuclear import receptor Kap95, we asked whether the transport of proteins with a nuclear localization sequence (NLS) was affected by ER stress. Indeed, nuclear import of newly synthesized NLS-mNeon was slowed (Figures 8F and S7B). In addition, some NLS-mNeon co-localized with Kap95 in cytosolic puncta (Figure S7B, C). By contrast, the nucleocytoplasmic distribution of BFP without an NLS was almost the same as in untreated cells, indicating that the observed import defect for NLS-mNeon was specific to active transport (Figure 8F). These results illustrate that DOMs uncovered unexpected localization changes not previously associated with ER stress.

## Discussion

We have established Dynamic Organellar Maps (DOMs) in yeast and combined them with fluorescence microscopy to explore protein abundance and localization changes during ER stress. The use of DOMs as a guide for subsequent imaging experiments enabled a systematic investigation and showed that protein relocalization is a major aspect of cellular reorganization during ER stress. Our results indicate that (1) many secretory pathway proteins, in particular cargos of the CPY pathway for vacuolar protein sorting, GPI-anchored proteins and Golgi-resident mannosyltransferases, accumulate in the ER, likely due to misfolding and subsequent ER retention; (2) some but not all lumenal ER proteins undergo reflux to the cytosol; (3) a subset of integral ER membrane proteins, including reticulon homology domain proteins, co-cluster and segregate from the remainder of the ER; (4) specific nucleoporins and importins form common cytosolic clusters. Interestingly, we observed that many stress-responsive proteins show either abundance or localization changes, but less frequently both. This tendency has been noted before (Litsios et al, 2024; Hein et al, 2025) and may reflect a more general regulatory principle.

Traditionally, microscopy and subcellular fractionation have been employed as parallel approaches to understand the spatial organization of cells. Since the advent of GFP, the imaging of fluorescently labeled proteins has become the predominant approach for analyzing protein localization in yeast (Huh et al, 2003; Chong et al, 2015; Kraus et al, 2017). However, recent advances in mass spectrometry, together with the realization that organelles need not necessarily be purified to determine their protein contents, have given rise to organellar mapping as another means of elucidating protein localization on a proteome-wide scale (Lundberg and Borner, 2019; Borner, 2020; Christopher et al, 2021). Indeed, our work indicates that the combination of DOMs and tailored follow-up microscopy studies offers several benefits.

First, DOMs detect native proteins, thus avoiding complications associated with tags such as GFP. For example, tail-anchored SNARE proteins, prenylated Rab GTPases, peroxisome proteins with classical targeting signals and ER lumenal proteins with retention signals all mislocalize when tagged at their C-termini. Furthermore, many proteins with signal sequences for entry into the ER or mitochondria cannot reach their destinations when tagged at their N-termini, and GPI-anchored proteins do not tolerate tags at either terminus. Accordingly, the five clear ER reflux cargos show incorrect localizations in the yeast strain collection with C-terminally GFP-tagged proteins, and the eleven ER-retained GPI-anchored proteins show incorrect localizations in both the C-and N-terminal GFP fusion collections (Huh et al, 2003; Weill et al, 2018; Meurer et al, 2018). Therefore, these proteins could not be found in earlier work on stress-induced protein relocalization (Breker et al, 2013). For many other proteins, it cannot be predicted a priori whether they retain their native localization after tagging (Weill et al, 2019).

Second, generating DOMs needs little equipment besides centrifuges and a mass spectrometer for label-free protein quantification. With our standard protocol, samples for up to four DOMs can be obtained in parallel by a single experimenter, and subsequent data analysis is aided by freely available software (https://domabc.bornerlab.org/QCtool; Schessner et al, 2023). Reproducibility between biological replicates was high and nearly 3000 proteins could be mapped across three experimental conditions. We here used mass spectrometry with data-dependent acquisition. However, data-independent acquisition can achieve a large increase in the number of mapped proteins (Schessner et al, 2023). Indeed, in preliminary measurements we found that yeast maps with 4000 proteins can be obtained with data-independent acquisition (unpublished results). This is similar to the number of proteins detectable by microscopy when tagged with GFP (Kraus et al, 2017).

Third, DOMs yield predictions for the localization of uncharacterized proteins. There was 93% agreement of our localization assignments from steady-state maps with the only previous organellar mapping study in yeast and 89% agreement with our literature-based reference database (Nightingale et al, 2019; Table S1D, E). Furthermore, DOMs provided high-confidence predictions for 190 proteins with previously ambiguous or unknown localization (Table S1F). However, it is important to bear in mind that many proteins have dual or multiple localizations (Lundberg and Borner, 2019), which the categorial single-organelle assignment for DOMs is not set up to accommodate. Therefore, new predictions need to be verified experimentally, and we emphasize that the main strength of DOMs is the detection of protein localization changes.

Various relocalizing proteins hint at new aspects of proteome remodeling during ER stress. For instance, cytosolic foci of nucleoporins have been found in yeast and metazoa under different stress conditions and may reflect the formation of protein condensates (Garcia et al., 2021; Thomas et al., 2023). Our systematic analysis uncovered redistribution of numerous nucleoporins belonging to specific nuclear pore subcomplexes. One possibility is that cytosolic nucleoporin foci result from pore complex assembly defects. Prolonged ER stress can overload the proteasome because of increased flux through the ER-associated degradation pathway and lead to misfolded protein accumulation and chaperone shortage in the cytosol (Schmidt et al, 2019). Therefore, the integration of newly synthesized nucleoporins into nuclear pore complexes could fail, due to either their own misfolding or the lack of other factors essential for assembly (Kuipers et al, 2022). Alternatively, cytosolic nucleoporin foci may arise from partial disassembly of pre-existing nuclear pore complexes. It has been proposed that the nucleoporin Nsp1, which forms pronounced foci upon DTT treatment, possesses chaperone activity and contributes to protein homeostasis in the cytosol (Otto et al, 2024). Interestingly, we find that Get3, another protein with conditional chaperone activity (Ulrich et al, 2022), shifts from the ER membrane towards the cytosol during stress. These observations invite the intriguing speculation that stress-induced relocalization of Nsp1 and Get3 serves to increase cytosolic chaperone capacity. Moreover, in our ER-centric experimental validation we did not follow up on several interesting predictions that concern other organelles (Table S5). For example, a number of proteins shifted away from mitochondria (Ccp1, Msp1, Ptc5, Ptc7, Tim9, Tim10, Tim13). These shifts could be caused by disturbed mitochondrial protein import or stress-induced reflux of correctly targeted proteins. Some spindle pole body proteins moved with the nuclear envelope (Mps3, Cdc31) but others did not (Spc42, Spc110, Nud1, Cnm67), perhaps hinting at reorganization of the spindle pole body as a result of stress-induced cell cycle arrest. Finally, a number of organelle contact site proteins were predicted to relocalize upon ER stress (Efr3, Epo1, Ist2, Num1, Osh6, Scs2, Swh1, Tcb1/2/3). The biological significance of these predictions awaits elucidation.

In conclusion, DOMs are an accessible and powerful technique for unbiased, comprehensive studies of protein localization changes in yeast. We therefore anticipate that DOMs are going to join the growing arsenal of techniques for systematic analyses that make yeast such an illuminating model organism.

## Materials and Methods

### Plasmids

Plasmids used in this study are listed in Table S7. To generate pFA6a-mScarlet-I3-HIS3 and pFA6a-mScarlet-I3-klTRP1, the mScarlet-I3 sequence (Gadella et al, 2023) was amplified from a synthetic DNA fragment and inserted into pFA6a-GFP(S65T)-HIS3 (Longtine et al, 1998) or pFA6a-mNeonGreen-klTRP1, replacing GFP or mNeonGreen. To generate pFA6a-HaloTag-klTRP1, the HaloTag7 sequence (Wilhelm et al, 2021) was amplified from a synthetic DNA fragement and inserted into pFA6a-mNeonGreen-klTRP1, replacing mNeonGreen. To generate pRS406-P_TEF_-TagBFP-Ubc6, the *GPD* promoter of pRS406-P_GPD_-mCherry-Ubc6 was replaced with the *TEF* promoter, creating pRS406-P_TEF_-mCherry-Ubc6. Subsequently, TagBFP was amplified from pRS303H-P_GPD_-TagBFP (Szoradi et al, 2018) and inserted into pRS406-P_TEF_-mCherry-Ubc6, replacing mCherry. To generate pFA6a-hph-P_GAL_-Kar2ss-sfGFP for N-terminal tagging of GPI-anchored proteins, the hygromycin resistance gene from pFA6a-hph was inserted into pFA6a-nat-P_GPD_-yeGFP (Janke et al, 2004), creating pFA6a-hph-P_GPD_-yeGFP. P_GPD_-yeGFP was then replaced by P_GAL_-Kar2ss-sfGFP amplified from pRS405-P_GAL_-Kar2ss-sfGFP-HDEL. To generate pNH605-P_ADH_-GEM-P_GAL_-NLS-mNeonGreen, the mNeonGreen sequence was amplified from pFA6a-mNeonGreen-HIS3 while adding an N-terminal SV40 nuclear localization sequence and inserted into pNH605-P_ADH_-GEM-P_GAL_ (Schmidt et al, 2019).

### Yeast strains

Strains used in this study were derived from *S. cerevisiae* W303 mating type a (SSY122) and are listed in Table S8. Gene tagging was done with PCR products of the plasmids listed in Table S7 (Longtine et al, 1998; Janke et al, 2004; Young et al, 2013).

### Subcellular fractionation

Yeast (SSY122) were cultured at 30°C in SCD medium containing 0.7% yeast nitrogen base (Merck), amino acids and 2% glucose. For steady-state maps, cultures were grown to early log phase (OD_600_ = 0.1-0.5) and 200 ODs of cells were harvested by centrifugation at 3,000 g at room temperature for 5 min. For treatment maps, cultures were grown to mid log phase and either (1) diluted to OD_600_ = 0.1, grown for 3 h and harvested, (2) diluted to OD_600_ = 0.15, grown for 1 h, treated with 8 mM DTT (Roche), grown for another 2 h and harvested, or (3) diluted to OD_600_ = 0.2, treated with 2 µg/ml tunicamycin (Merck), grown for another 3 h and harvested. Cells were resuspended to 20 ODs/ml in 10 ml spheroplast buffer (50 mM Tris-HCl pH 7.5, 1 M sorbitol, 0.5 mM MgCl_2_). An aliquot of the cell suspension was diluted 1:40 in water and the OD_600_ was measured, which gave a reading of about 0.5. To digest the cell wall, 40 µl zymolyase T100 (Biomol) were added to the cell suspension and cells were incubated at 30°C for up to 20 min. During this time, spheroplast formation was monitored by periodically measuring the OD_600_ as described above. Cell wall removal renders cells osmotically sensitive, leading to cell rupture and a drop in OD_600_ when they are diluted in water. Once the OD_600_ had fallen below 0.15, cell wall digestion was terminated by transferring the cell suspension onto ice. All subsequent steps were carried out in the cold. Spheroplasts were washed three times by pelleting at 1,000 g for 2 min and gentle resuspension in 5 ml spheroplast buffer. After the last wash, cells were resuspended in 3.5 ml hypo-osmotic lysis buffer (25 mM Tris-HCl pH 7.5, 200 mM sorbitol, 1 mM EGTA, 0.5 mM MgCl_2_, Roche protease inhibitors without EDTA) and sample volumes were adjusted to 4 ml with lysis buffer. Lysed spheroplasts were homogenized by 20 strokes in a Dounce homogenizer with a tight-fitting pestle (Thermo Fisher Scientific, catalog number 8853000007). Of the resulting homogenate, 50 µl were collected for full proteome analysis and combined with 12.5 µl 5x SDS resuspension buffer (50 mM Tris-HCl pH 8.1, 12.5% w/v SDS). DTT was added to a final concentration of 0.1 mM to prevent oxidation, samples were frozen in liquid nitrogen and stored at-80°C. The remainder of the homogenate was subjected to a clearing spin at 300 g at 4°C for 5 min to remove unbroken cells. The supernatant was transferred into a 15 ml conical tube and centrifuged at 1,000 g at 4°C for 10 min. The resulting 1k pellet was kept on ice, the supernatant was transferred into a new 15 ml conical tube and centrifuged at 3,000 g at 4°C for 10 min. The resulting 3k pellet was kept on ice, the supernatant was transferred into an ultracentrifuge tube and centrifuged with a TLA-110 rotor in an Optima TLX tabletop ultracentrifuge (Beckman) at 6,000 g at 4°C for 15 min. The resulting 6k pellet was kept on ice, the supernatant was transferred into a new ultracentrifuge tube and the procedure was repeated to obtain 12k, 24k and 78k pellets by successive centrifugation steps at 12,000 g for 20 min, 24,000 g for 20 min and 78,000 g for 30 min. Of the 78k supernatant, 800 µl were mixed with 200 µl 5x SDS resuspension buffer (see above). All pellets were taken up in 1x SDS resuspension buffer. To ensure protein concentrations of at least 1 µg/µl, 200 µl 1x SDS resuspension buffer were used for the 1k pellet, 100 µl for the 3k, 6k, 12k and 24k pellets, and 150 µl for the 78k pellet. All samples were incubated at 65°C for 3 min. The 1k pellet was further solubilized by 15 cycles of 30 s on/off at maximum intensity with a Bioruptor (Diagenode). DTT was added to all samples to a final concentration of 0.1 mM and samples were frozen in liquid nitrogen and stored at-80°C.

### Peptide generation for mass spectrometry

Proteins from total proteome, pellet and 78k supernatant samples were converted into peptides for mass spectrometry as described (Itzhak et al, 2019). Briefly, protein concentrations were determined with a BCA assay (Pierce) and samples were diluted to 1 µg/µl with 1x SDS resuspension buffer. Proteins were precipitated with ice-cold acetone, resuspended in digestion buffer containing 8 M urea and 1 mM DTT, alkylated with 5 mM iodoacetamide and digested with LysC and trypsin. Peptides were acidified with trifluoroacetic acid, extracted with SDB-RPS Stop and Go Extraction (Stage) tips and eluted with 1.25% v/v ammonium hydroxide in 80% v/v acetonitrile. Sample volume was reduced to less than 5 µl with a vacuum Concentrator plus (Eppendorf) and peptides were resuspended in solution A (0.1% trifluoroacetic acid, 2% acetonitrile). Concentrations were determined with a NanoDrop 1000 spectrophotometer (Thermo Fisher Scientific) and adjusted to 150 ng/µl. Peptides were frozen in liquid nitrogen and stored at-80°C.

### Mass spectrometry

Mass spectrometric measurements were done with data-dependent acquisition (Schessner et al, 2023). Nanoflow reversed-phase chromatography was performed with an EASY-nLC 1200 ultra-high-pressure system coupled to an Orbitrap Exploris 480 mass spectrometer via a nano-electrospray ion source (Thermo Fisher Scientific). A binary buffer system with mobile phases A (0.1% v/v formic acid) and B (0.1% v/v formic acid, 80% acetonitrile) was used. Peptides were separated in 110 min on a 50 cm × 75 µm (i.d.) column, packed in-house with ReproSil-Pur C18-AQ 1.9 µm silica beads (Dr. Maisch GmbH). The column was operated at 60°C. Purified peptides were loaded onto the column in phase A and eluted with a linear 5-30% gradient of phase B, followed by washout and column re-equilibration. The mass spectrometer was controlled by Xcalibur software (Thermo Fisher Scientific) and operated in top 15 scan mode with a full scan range of 300-1650 Th. Survey scans were acquired at 60,000 resolution, with automatic gain control (AGC) set to 300% and a maximum ion injection time of 25 ms. Charge states were filtered for 2-5. Precursor ions were isolated in a window of 1.4 Th, fragmented by higher-energy collisional dissociation with normalized collision energies of 30%. Fragment scans were performed at 15,000 resolution with a maximum injection time of 28 ms, AGC set to 100% and a dynamic precursor exclusion for 30 s.

### Mass spectrometry raw data analysis

For protein identification, mass spectrometry raw data were analyzed in MaxQuant version 2.0.1.0 (Tyanova et al, 2016a). Match between runs and label-free quantification (LFQ) were enabled. The minimum LFQ count was set to 1. Default parameters were used for all other settings. The MaxQuant experimental design restricted matching to equivalent and adjacent fractions within the six-fraction profiling speed gradient. Cytosol and full proteome fractions were only matched to equivalent fractions. For ER stress maps, matching was only possible within a treatment condition. Spectra were searched against the SwissProt FASTA Saccharomyces cerevisiae database downloaded from UniProt (6,750 entries). The resulting intensity data and normalized profiles are provided as Supplemental Datasets 1 and 2. Mass spectrometry data generated here will be made available at the ProteomeXchange Consortium via the PRIDE partner repository.

### Generation of compartment marker list

A reference database was created containing 5389 proteins for which abundance estimates in molecules per cell were available (Table S2A; Ho et al, 2018). Eighteen localization categories were defined: actin-associated, cell wall, COPI coat, COPII coat, cytosol, ER, endosomes, ER-Golgi intermediate compartment (ERGIC), Golgi, lipid droplets, mitochondria, nucleus, nuclear envelope, peroxisomes, plasma membrane, proteasome core, ribosome core and vacuole. Two additional categories were ‘ambiguous’ (proteins for which evidence suggested localization to more than one subcellular compartment) and ‘unknown’ (proteins for which no assignment was possible). Based on an analysis of the GFP fusion collection (Kraus et al, 2017; data downloaded from the CYCLoPs web site), the Saccharomyces Genome Database and primary literature, proteins were manually assigned to one of the above categories.

The 2971 protein groups present in steady-state maps of unperturbed yeast were taken from the reference database and used for iterative training of the SVM module in DOM-ABC (see below). The first training set consisted of 868 protein groups, which were the 100 most abundant protein groups in the categories cytosol, ER, mitochondria, nucleus, plasma membrane and vacuole including endosomes, as well as all available protein groups in the categories Golgi, lipid droplets, nuclear envelope, peroxisomes, proteasome core and ribosome core. Proteins from the remaining categories were omitted because their localizations are inherently ambiguous. COPI, COPII and ERGIC proteins show complex distributions at the ER/Golgi interface and proteins related to the actin cytoskeleton are cytosolic but can associate with the plasma membrane. In addition, cell wall proteins were omitted because the cell wall had to be removed as part of the subcellular fractionation procedure. A first round of training achieved 94% recall, i.e. the SVM prediction agreed with the reference database for 815 protein groups. Furthermore, there were 682 additional protein groups for which a localization was predicted with at least medium confidence and matched the annotation in the reference database. A second training set was assembled by combining the 815 correctly recalled protein groups and the 682 additional protein groups correctly predicted during the first training round. The second round of training achieved 99.9% recall and correctly predicted the localization of 269 additional protein groups, and these results were used to assemble the third training set. This procedure was repeated until, after the seventh iteration, recall of the 1908 protein groups in the training set was 100% and no further protein groups were predicted correctly with at least medium confidence. The resulting final compartment marker set consisted of 1908 protein groups, corresponding to 1937 unique proteins.

### Organellar mapping data analysis in DOM-ABC

The data required to recapitulate the DOM-ABC analyses are provided as Supplemental Dataset 3. These include trimmed protein groups files (steady-state maps, ER stress maps combined, ER stress maps single) as input, the custom yeast compartment marker list, the Uniprot tab used for gene annotation, the DOM-ABC settings and output.json files, and instructions for processing. Data filtering, annotation, normalization, quality control, and mapping analyses were performed as described (Schessner et al, 2023). Briefly, the protein groups output file from MaxQuant is loaded into the online tool domaps version 1.0 (https://github.com/JuliaS92/SpatialProteomicsQC/tree/1.0), which formats the data for downstream analysis and quality control. Intensities in each of the six organelle fractions (1,000-78,000 x g pellets) are normalized to the total summed intensity across all fractions to obtain 0-to-1 normalized profiles. These can be directly compared between proteins, irrespective of relative abundance, and reflect subcellular distribution. Normalized profiles are then used for all downstream analyses, including calculation of map depth, fraction correlation evaluation, visualization by PCA, protein shift analysis, and compartment classification by SVMs. Relevant parameters were:

(1) Data annotation. Proteins were annotated with the compartment markers list and current gene names obtained from the Saccharomyces Genome Database.

(2) Data filtering. Only profiles with at least three consecutive fraction MS intensities and a minimum average MS count of two were retained.

(3) Principal component analysis (PCA). The ‘fix aspect ratio by variability’ was deselected to optimize axis scaling for visualization. PC1 versus PC2 were plotted as the most informative principal components to show map resolution. For steady-state maps, all six replicates were combined into one dataset and processed jointly in the ‘analysis’ module of DOM-ABC. For ER stress maps, data for each condition were separated into three protein groups files and processed individually to generate three.json files. These were then loaded into the benchmark module and jointly subjected to PCA. To generate the PCA plots of individual maps, the joint data were reloaded into the ‘analysis’ module.

(4) SVM compartment classification. Default settings were used to train and benchmark SVMs (C parameter range 1-30, gamma parameter range 1-50, 5 iterations with built-in five-fold cross validation). For model training, all markers were used and the optimized parameters were used for classification (test set proportion set to 0). For performance benchmarking, markers were split 80:20 into training and leave-out test sets (test set proportion set to 0.2). Models were trained with five-fold cross-validation on the training set only. Optimized SVMs were then applied to the test set to evaluate prediction performance via the F1 score (i.e. the harmonic mean of recall and precision). Average F1 scores across 20 sub-samples (i.e. 75%) of the predictions on the test set were then calculated for each organelle. The classes ‘lipid droplets’ and ‘peroxisomes’ had too few members to generate informative test sets and were excluded from performance benchmarking. For steady-state maps, all six replicates were combined into one dataset and processed jointly in the ‘analysis’ module of DOM-ABC to generate a single.json file. This file was then loaded into the ‘benchmark’ module of DOM-ABC for SVM classification. For ER stress maps, data for each condition were separated into three protein groups files and processed individually in the analysis module to generate three.json files. These were then loaded into the benchmark module and jointly subjected to SVM analysis. The output of the SVM classification is a probability score for the likelihood of model fit for each of the 12 compartments represented in the compartment marker list (see above). For each protein, scores across compartments therefore add up to 1. Protein are assigned to the compartment with the highest score. Assignments are grouped into confidence classes depending on the magnitude of the score: >0.95 = very high, >0.8 = high, >0.65 = medium, >0.4 = low, <0.4 = best guess. For Figure 1C, predictions with ‘very high’ and ‘high’ confidence were further grouped as ‘high confidence’, and predictions with ‘low confidence’ and ‘best guess’ as ‘low confidence’.

(5) Movement-reproducibility (MR) analysis. To detect proteins with localization changes, default settings were applied for data pre-filtering (cosine correlation >0.9) and statistical testing (static data proportion = 0.75, number of iterations = 11). For each protein, the profiles from control cells are subtracted from profiles from DTT-or tunicamycin-treated cells to obtain a ‘delta profile’. Proteins without significant changes have delta profiles close to baseline. To identify significantly deviating delta profiles, a robust multivariate outlier test is performed. P-values from three replicates are then combined with the Fisher method. The joint p-value is corrected with the Benjamini-Hochberg method and-log10 transformed to obtain the movement (M) score. M = 2 means that a protein undergoes a significant movement with an estimated FDR of 1%. Here, an M-score of 1.3 was chosen as cut-off for significance (FDR <5%). As an additional stringency filter, the direction of movement had to be consistent across replicates. The test therefore calculates the pairwise Pearson correlation of delta profile replicates (Rep1 vs Rep2, Rep1 vs Rep3 and Rep2 vs Rep3). The median of these three values was chosen as the R-score, with a value of 0.8 as cut-off for reproducibility. Finally, proteins had to have a p-value for movement <0.05 in at least two of the three replicates to qualify as hits. DOM-ABC performs a data quality filtering step prior to MR analysis so that only profiles with high replicate reproducibility (all pairwise cosine correlations >0.9) and only proteins profiled in all three replicates of both compared conditions are included. Hence, the number of proteins in the MR analysis is usually lower than the number of mapped proteins.

### Full proteome quantification analysis

Full proteome LFQ intensities were extracted from MaxQuant protein groups files and analyzed with Perseus software V1.6.2.3 (Tyanova et al., 2016b). Proteins were filtered to remove reverse hits, proteins ‘only identified by site’ and potential contaminants.

For pairwise ‘volcano’ analyses, proteins were required to include at least three measured datapoints within one condition. Following log transformation, missing data were imputed from a normal distribution with a downshift of 1.8 standard deviations and a width of 0.3 standard deviations. Data were then analyzed with a two-tailed t-test to identify proteins with altered abundance. Non-linear significance cut-offs (i.e. the ‘volcano lines’) were defined via Perseus’ permutation-based FDR calculation (FDR <5%, S0 = 0.1).

For 1D-annotation enrichment, proteins were annotated with GO terms in Perseus and the results of the volcano analyses were analyzed with default settings.

For the full proteome PCA plot (Figure S3A), proteins from all three conditions were analyzed jointly. Proteins were required to include at least three measured datapoints within one condition. Following log transformation, missing data were imputed from a normal distribution with default settings. Data were then subjected to PCA.

For copy number and concentration estimates, the ‘Proteomic Ruler’ method was applied as a plugin in Perseus 2.0.11.0 (Yu et al, 2020). Raw intensities of the six full proteomes of unperturbed cells were filtered to remove reverse hits, proteins ‘only identified by site’ and potential contaminants, and annotated with average molecular weights from the yeast FASTA sequence database used for MaxQuant processing. The proteomic ruler tool was then applied using average molecular weights to normalize intensities, processing all six data columns separately. An estimate for ploidy was obtained from entries 109469, 108315 and 108196 of the BioNumbers database (Milo et al, 2010), which yielded an average of 0.022 pg DNA/cell. Assuming 12.1 million base pairs or 0.013 pg per yeast genome, 0.022 pg DNA/cell correspond to 1.7 genomes/cell. An estimate for intracellular protein concentration was obtained from BioNumbers entries 100427, 100430, 100452, 111978, 109469, 108315, 106225, 100490, which yielded 94 g/l for haploid and 79 g/l for diploid yeast. Interpolation to a ploidy of 1.7 gave an estimated concentration of 83 g/l.

### Cytosolic pool analysis

To estimate protein cytosolic pools, a second set of abundance profiles was generated in DOM-ABC that included the six organellar fractions and also the cytosol fraction. Normalized intensities in each fraction were weighted with the corresponding relative protein yields as measured by BCA assay. Weights for each fraction were calculated as average percentual protein recovery across replicates. Weighted intensities were divided by the sum across all fractions to obtain 0-to-1 normalized weighted intensities. These seven-datapoint profiles reflect percentage recovery of a protein across subcellular fractions. The cytosol fraction corresponds to the actual cytosolic pool (which is excluded from the six-datapoint profiles used to generate organellar maps, see above). The sum of the first six fractions indicates a protein’s non-cytosolic pool.

For steady-state maps, proteins were annotated with cytosolic pools to identify proteins with potential dual cytosolic and non-cytosolic localizations (Table S1C) and determine the extent of organellar leakage during fractionation (Figure S1). For ER stress maps, cytosolic pools were analyzed to identify proteins that shift to or away from the cytosol upon ER stress. First, proteins not quantified in all maps were excluded. Second, proteins minimally had to have a measured cytosolic pool in all three replicates of one treatment condition. For the remaining 2204 proteins, the cytosolic pool change was analyzed in control relative to DTT-or tunicamycin-treated cells with a paired two-tailed t-test. Since generation of the cytosolic pool data required several processing steps, which may contribute additional noise, the results for DTT-and tunicamycin-treated cells were combined to increase stringency and statistical power. The two individual p-values were combined with the Fisher method and the joint p-values were corrected for multiple testing with the Benjamini-Hochberg method. To qualify as hits, proteins had to have: an overall FDR <5%; significant individual p-values with both DTT and tunicamycin treatment (<0.05 for the DTT set and <0.1 for the slightly noisier tunicamycin set); an absolute cytosolic pool change >10% with DTT and tunicamycin treatment; the same direction of change (i.e. towards or away from the cytosol) under both conditions. 74 proteins passed these filters (Table S4C). Changes in the cytosolic pool are shown only for the DTT data in Figure 5A, but the identification of significant hits was performed as describe here.

### Comparison of SVM localization predictions with reference database

The SVM predictions included 12 annotated subcellular localizations, the reference database 20. To enable a direct comparison of the localization predictions, the reference database categories ‘endosomes’ and ‘vacuole’ were pooled (to just ‘vacuole’), and also the categories ‘cell wall’ and ‘plasma membrane’ (to just ‘plasma membrane’). Proteins with categories not represented in the SVMs were removed (‘actin-associated’, ‘ambiguous’, ‘COPI’, ‘COPII’, ‘ERGIC’, ‘unknown’). This left 2554 proteins with matching categories. Furthermore, proteins were annotated with the cytosolic pools determined in this study. Proteins with organelle assignments and an additional cytosolic pool >30% were considered to have a dual localization (Table S1C).

### Identification of proteins shifting towards the ER during ER stress

First, profile Pearson correlation of mapped proteins with the average ER marker profile was determined for control and DTT-treated cells. For each protein, the change of correlation with ER markers was calculated as Delta correlER = (correlationDTT ER) – (correlationCon ER). A positive Delta CorrelER identified proteins that correlated better with ER markers under ER stress. To identify proteins shifting towards the ER, the following filters were applied: hit in the DTT MR analysis (M >1.3, R >0.8); positive Delta CorrelER; Pearson correlation with average ER marker profile in DTT-treated cells >0.75; protein has post-ER secretory pathway localization in untreated cells according to reference database or SVM classification. This analysis identified 86 proteins. Inspection of the 410 DTT hits identified three proteins (Ape3, Sln1, Toh1) that narrowly failed the second filter but passed all other filters and had a convincing shift towards the ER in PCA plots. They were therefore added to the final list of 89 proteins (Table S5). For three of these (Dcr2, Lcb5, Epo1), UniProt information did not predict transmembrane domains or ER-targeting signal peptides, suggesting that they are peripheral membrane proteins.

### Identification of ER lumenal proteins

Proteins in the reference database (Table S2A) were filtered for ER localization. These 315 proteins were annotated with UniProt information on transmembrane domains and ER-targeting signal peptides. The 17 ER proteins with a signal peptide and no transmembrane domains were considered lumenal. Based on the localization data generated in this study, we manually added to this list two proteins (Fpr2, Fmp27) from the pool of proteins with ‘ambiguous’ localization in the reference database.

### Identification of post-ER secretory pathway proteins

The 2320 proteins available for MR analysis in DTT-treated versus control cells (Table S4A) were annotated with UniProt information on transmembrane domains, ER-targeting signal peptides and mitochondria-targeting transit peptides. Proteins were filtered to contain an ER-targeting signal peptide and/or one or more transmembrane domains but did not contain a mitochondrial-targeting transfer peptide. Remaining proteins were filtered for a predicted localization to post-ER secretory compartments. For this, we principally used the reference database but manually augmented it with data from this study. This procedure identified 243 post-ER secretory pathway proteins present in the DTT profiling set (Table S4E).

### Comparison of SVM localization predictions with previous yeast data

The roughly 900 localization predictions in Nightingale et al, 2019 were category-matched to the SVM predictions generated here, where possible, and matched up via protein IDs. Localization predictions for the 775 proteins that could thus be compared across both datasets showed 93% agreement (Table S1E).

### Microscopy

Cells were cultured as described above and treated with 8 mM DTT for 2 h where indicated. Cells expressing GFP-tagged GPI-anchored proteins under an estradiol-inducible promoter system were cultured in the presence of 50 nM estradiol (Sigma) for 16 h prior and DTT treatment was done in SCD containing 50 nM estradiol and 50 mM HEPES pH 7.5. HEPES was included to avoid quenching of extracellularly exposed GFP by the low pH of SCD medium. Cells from 1 ml culture were harvested by centrifugation at 10,000 g at room temperature for 2 min. The supernatant was removed and cells were resuspended in 20 µl medium. For staining with the silicon-rhodamine HaloTag ligand (Lukinavicius et al, 2013), cells were harvested, resuspended in 30 µl PBS containing 500 nM HaloTag ligand and incubated at 800 rpm at room temperature for 10 min. Three µl cell suspension were mounted on coverslips and covered with 1% w/v agarose pads made with SCD medium. For imaging of GPI-anchored proteins, agarose pads were made with PBS instead of SCD to avoid quenching of GFP fluorescence.

Images were acquired with a Nikon Ti2-W1 spinning disk confocal microscope equipped with a Yokogama W1 scanhead, a Nikon Plan Apo 100x/NA 1.45 objective and a Zyla 4.2P CMOS camera (Figures 5C, 5E, 8B, 8E, S6A, S7C), or with a Nikon Ti2 widefield microscope equipped with a Nikon Plan Apo 100x/NA 1.45 objective and a Hamamatsu Orca Fusion-BT camera (in Figures 6C, 7B, 7D, S6B-D, S7B). Images for Figures 5E, 6C, 7B, 7D and S6B-D were acquired as Z-stacks with a step size of 200 nm to facilitate subsequent image deconvolution. For the systematic analysis of GFP-tagged nucleoporins (Table S6), cells were treated and imaged by widefield microscopy as above.

Images were anonymized with the “Blind Analysis Tools” plugin in ImageJ (https://imagej.net/plugins/blind-analysis-tools) before visual analysis to prevent user bias. Puncta formation was assessed independently by two individuals. The resulting rare cases of disagreement were resolved by a second round of joint assessment.

Confocal images for Figures 5C, 8B, 8E, S6A and S7C were processed with Fiji software. General background was subtracted in all channels with the rolling ball algorithm (radius = 150 pixels, or 9.75 µm), regions of interest were selected and brightness/contrast was adjusted once in each channel with the ‘auto’ command. Widefield images for Figure S7B were processed the same except that no auto-adjustment was applied to the NLS-mNeon signal. Confocal images for Figure 5E and widefield images for Figures 6C, 7B, 7D and S6B-D were deconvolved with the Richardsson-Lucy algorithm (30 iterations, noise level set to automatic, background subtraction activated) in NIS Elements software (Nikon).

### Nuclear import assay

Yeast (SSY4574) were cultured as described above. Control cells were diluted to OD_600_ = 0.15 and grown for 1 h. Cells to be exposed to ER stress were diluted to OD_600_ = 0.25, treated with 8 mM DTT and grown for 1 h. Each culture was split in two, one new culture received no further treatment and was used to determine background autofluorescence in the GFP channel, the other new culture was treated with 500 nM estradiol for 50 min to induce expression of NLS-mNeonGreen. Cells were imaged by widefield microscopy as described above and single optical sections in z were acquired.

Image quantification was done with Fiji software. Background signal was subtracted in all channels with the rolling ball algorithm as above and a cell mask was generated based on the combined BFP and brightfield images using global automatic pixel intensity thresholding followed by watershed segmentation. A nucleus mask was generated the same way based on the Pus1-mScarlet image. Cell and nucleus masks were used to measure the areas, mean BFP and mean mNeonGreen pixel intensities of individual cells and nuclei. Cell measurements were done for objects with a size of 9-27 µm^2^ and a circularity of 0.5-1, nucleus measurements for objects with a size of 1.1-7 µm^2^ and a circularity of 0.1-1. Nucleus measurements were assigned to the corresponding cell measurements. Cell measurements with no or more than one nucleus measurement were removed. To determine nuclear enrichment of NLS-mNeonGreen, mean cell and nuclear mNeonGreen pixel intensities were first determined from cells not treated with estradiol (>250 cells per condition and replicate). These background fluorescence values were substracted from the mean cell and nuclear pixel intensities in estradiol-treated cells, which expressed NLS-mNeonGreen. Cells with background-corrected mean pixel intensities of less than 1 were removed. Nuclear enrichment in the remaining cells was then calculated by dividing the background-corrected mean pixel intensity of each nucleus by the background-corrected mean pixel intensity of the corresponding cell (>200 cells per condition and replicate). Autofluorescence in the BFP channel was negligible, so that nuclear enrichment of BFP was calculated without background correction by dividing the mean pixel intensity of each nucleus by the mean pixel intensity of the corresponding cell (>250 cells per condition and replicate). Data from three biological replicates were combined, yielding >800 cells analyzed per condition.

## Supporting information

Supplemental Database 1

Supplemental Database 2

## Acknowledgements

We thank Alexandra Davies for help with mass spectrometry, Kai Johnsson for the SiR HaloTag ligand, the Nikon Imaging Center at Heidelberg University for assistance, and Anne Schlaitz and all Schookees for incisive comments on the manuscript. This work was supported by grant SCHU 2364/3-1 from the German Research Foundation (DFG) to SS. GHHB would like to thank Matthias Mann for his continued support. The authors gratefully acknowledge the data storage service SDS@hd supported by the Ministry of Science, Research and the Arts Baden-Württemberg (MWK) and the DFG through grant INST 35/1503-1 FUGG. The authors declare no competing financial interests.

## Author contributions

Conceptualization: Georg Borner, Anna Platzek, Sebastian Schuck; Investigation: Anna Platzek, Klara Odehnalova, Julia Schessner; Formal analysis: Georg Borner; Funding acquisition and supervision: Sebastian Schuck; Writing – original draft: Georg Borner, Anna Platzek, Sebastian Schuck; Writing – review and editing: all authors.

---

**Figure S1.**
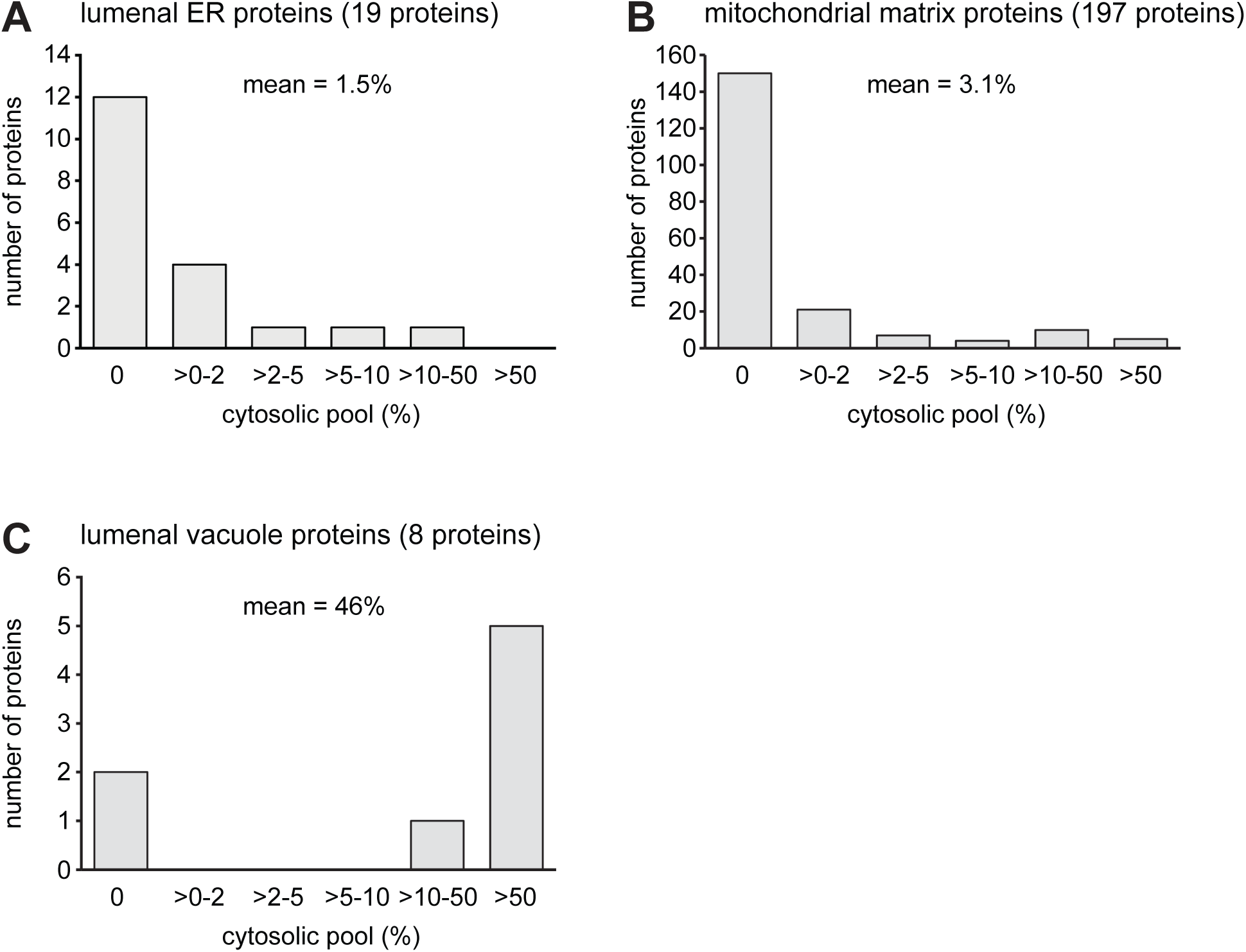
Leakage of lumenal organelle proteins from the ER, mitochondria and the vacuole into the cytosol. **(A)** Number of lumenal ER proteins with cytosolic pools of 0, >0-2, >2-5, >5-10, >10-50 or >50%. Lumenal proteins were predicted based on the presence of a signal peptide and the absence of a transmembrane domain. Cytosolic pool estimates were derived by dividing protein intensities in the cytosol fraction by the summed intensities in the cytosol and all organelle fractions (Table S1B). The mean cytosolic pool of lumenal ER proteins was 1.5%, indicating that ER membranes resealed quickly after cell lysis. **(B)** As in panel A but for soluble mitochondrial proteins that contain transit peptides, do not have transmembrane domains and have not been identified as intermembrane space proteins (Vögtle et al, 2012). These proteins are predicted to localize to the mitochondrial matrix. Their mean cytosolic pool is 3.1%, indicating that mitochondria remain intact during cell lysis or reseal quickly. **(C)** As in panel A but for lumenal vacuole proteins. Their mean cytosolic pool is 46%, indicating that many vacuoles are broken during cell lysis, leading to extensive leakage of lumenal proteins.

**Figure S2.**
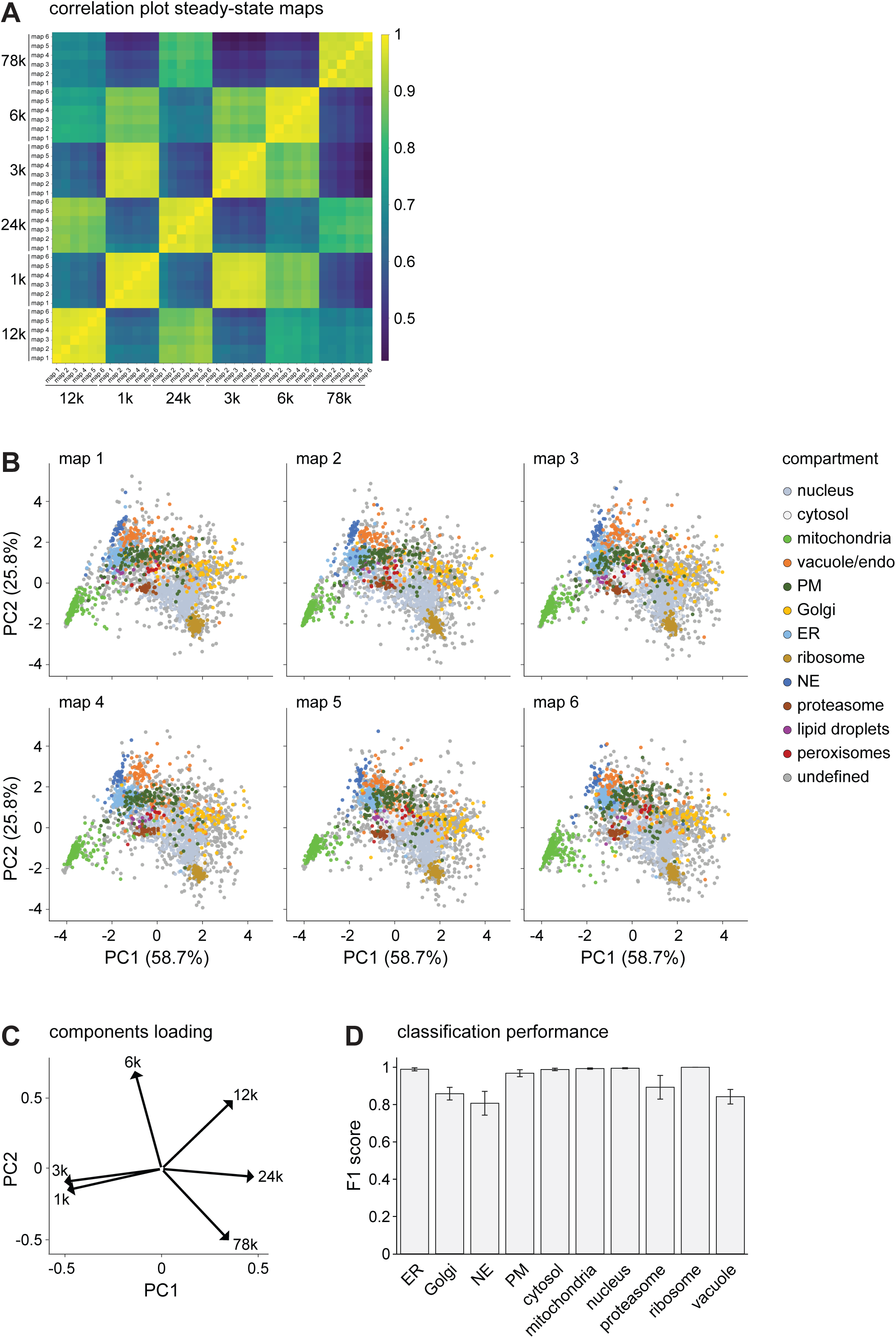
Quality assessment of organellar maps of unperturbed yeast. **(A)** Heatmap correlation plot of steady-state maps. Plotted are the pairwise Pearson correlations of all subcellular fractions and replicates. Fraction replicates have high reproducibility (R > 0.94). **(B)** PCA plots of all six steady-state organellar maps obtained from untreated yeast. Pre-defined compartment markers are colored, all other proteins are classified as’undefined’. The shared proteins of these maps were used to generate the map shown in Figure 1B. **(C)** PCA loadings plot for steady-state maps. The plot shows to what extent abundance in each subcellular fraction contributes to the position of a protein in the PCA plot shown in panel B. **(D)** Performance assessment of SVM classifications for steady-state maps. F1 scores indicate classification performance for each compartment, with perfect recall and precision yielding a score of 1. Error bars show standard deviation of repeated subsampling of predictions on the test set. Lipid droplets and peroxisomes had less than 20 marker proteins and performance for these comparments was not evaluated. NE, nuclear envelope; PM, plasma membrane.

**Figure S3.**
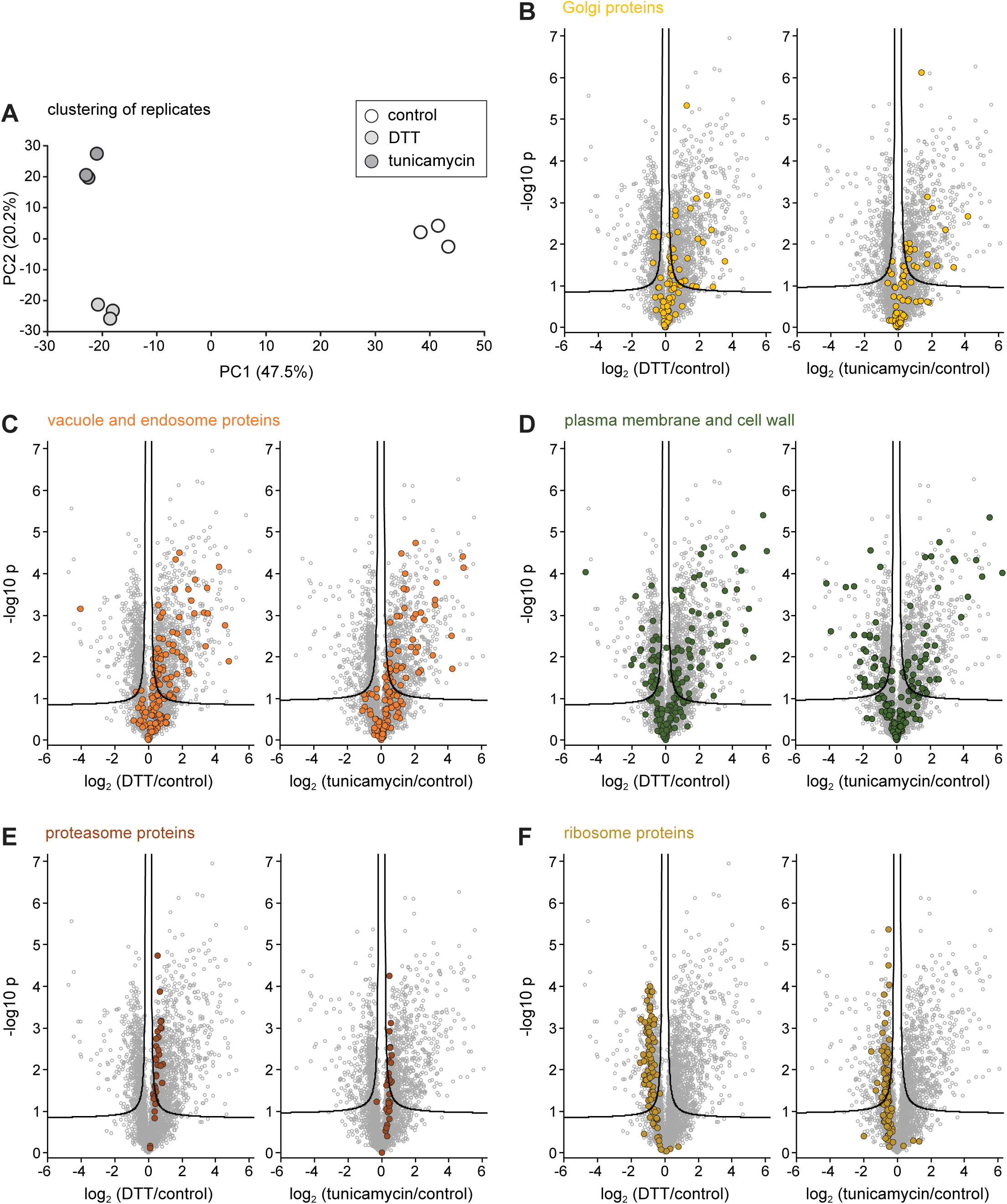
Protein abundance changes upon ER stress. **(A)** PCA plot of all three treatments and replicates based on protein intensities from full proteome samples. The plot shows that replicates cluster tightly within conditions and that all three conditions are clearly resolved. **(B)** Volcanos plots of full proteomes, showing abundance changes upon DTT or tunicamycin treatment. Plotted are the log_2_ fold changes upon treatment. P-values for the significance of changes were calculated with a two-sided t-test (n = 3). Volcano lines indicate 5% false discovery rate cut-offs based on data permutation. Golgi proteins are highlighted. **(C)** As in panel B but for vacuole and endosome proteins. **(D)** As in panel B but for plasma membrane and cell wall proteins. **(E)** As in panel B but for structural proteins of the proteasome. **(F)** As in panel B but for structural proteins of the ribosome.

**Figure S4.**
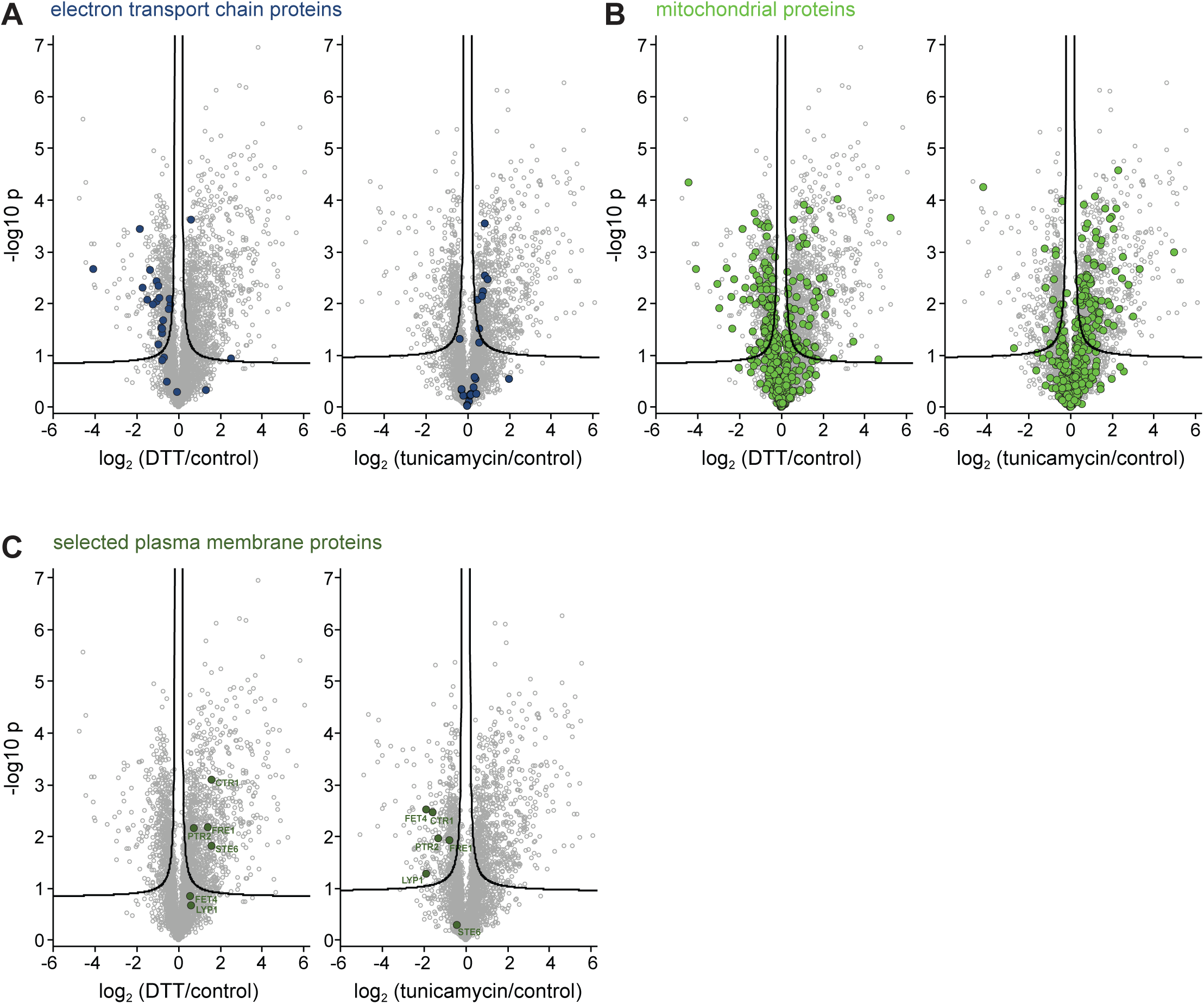
Divergent protein abundance changes upon DTT and tunicamycin treatment. **(A)** Volcanos plots of full proteomes, showing abundance changes upon DTT or tunicamycin treatment. Plotted are the log_2_ fold changes upon treatment. P-values for the significance of changes were calculated with a two-sided t-test (n = 3). Volcano lines indicate 5% false discovery cut-offs based on data permutation. Components of the mitochondrial electron transport chain are highlighted. DTT caused a decrease in the levels of most components whereas tunicamycin caused an increase. **(B)** As in panel A but mitochondrial proteins are highlighted. DTT tended to cause a downregulation of mitochondrial proteins and tunicamycin an upregulation. **(C)** As in panel A but selected plasma membrane proteins are highlighted that are upregulated upon DTT treatment and downregulated upon tunicamycin treatment. All of the highlighted proteins are involved in metabolite transport across the plasma membrane.

**Figure S5.**
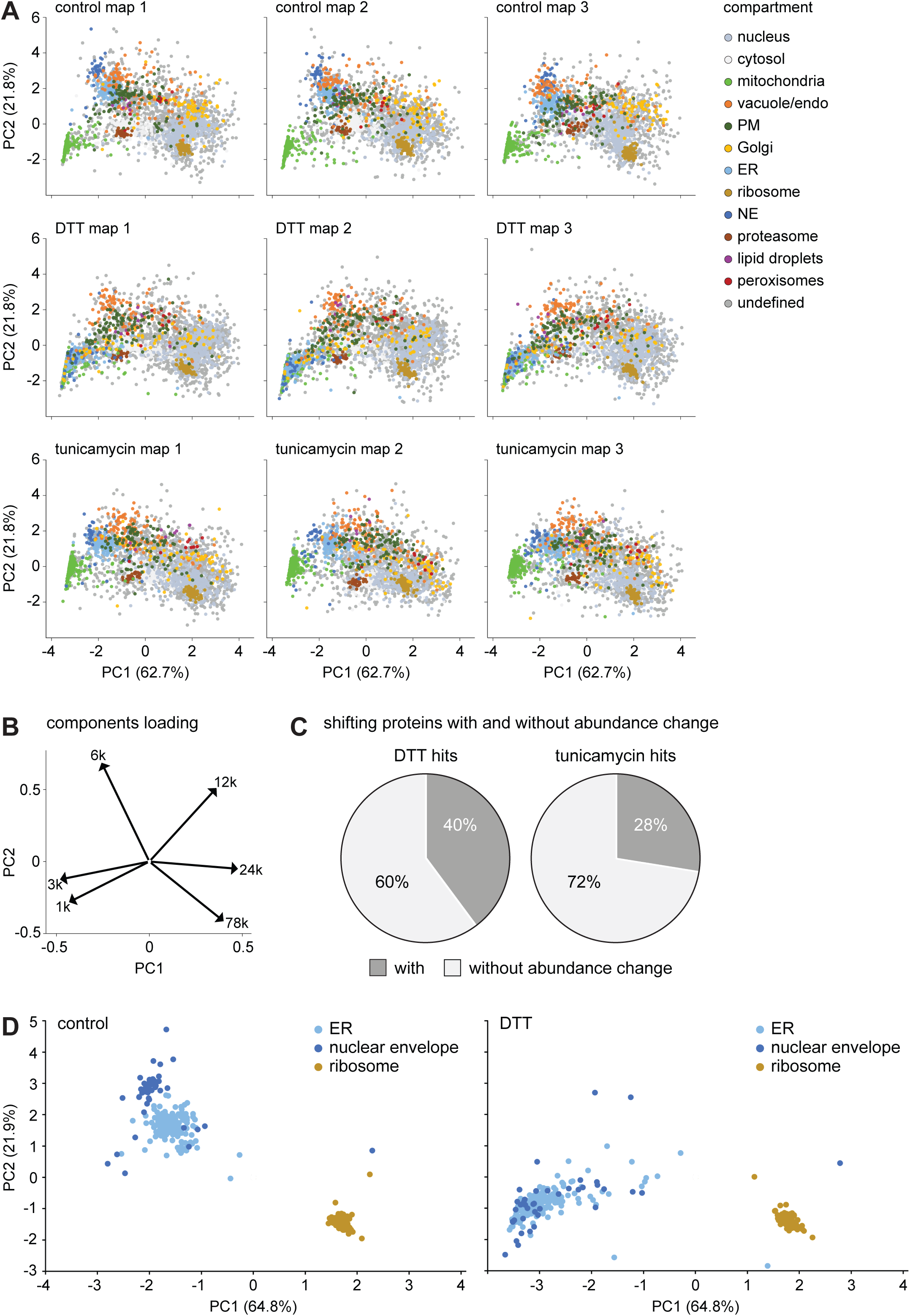
Protein localization changes upon ER stress. **(A)** PCA plots of all organellar maps obtained from control, DTT-treated and tunicamycin-treated cells. Pre-defined compartment markers are colored, all other proteins are classified as’undefined’. The averaged individual profiles of each condition were used to generate the maps shown in Figure 3A. **(B)** PCA loadings plot. The plot shows to what extent abundance in each subcellular fraction contributes to the position of a protein in the PCA plots in panel A. **(C)** Fraction of shifting proteins with and without accompanying abundance change. The majority of moving proteins do not change in abundance. **(D)** PCA plots of control and DTT maps highlighting the shifts of the ER and nuclear envelope clusters upon DTT treatment. The ribosome cluster is shown for reference.

**Figure S6.**
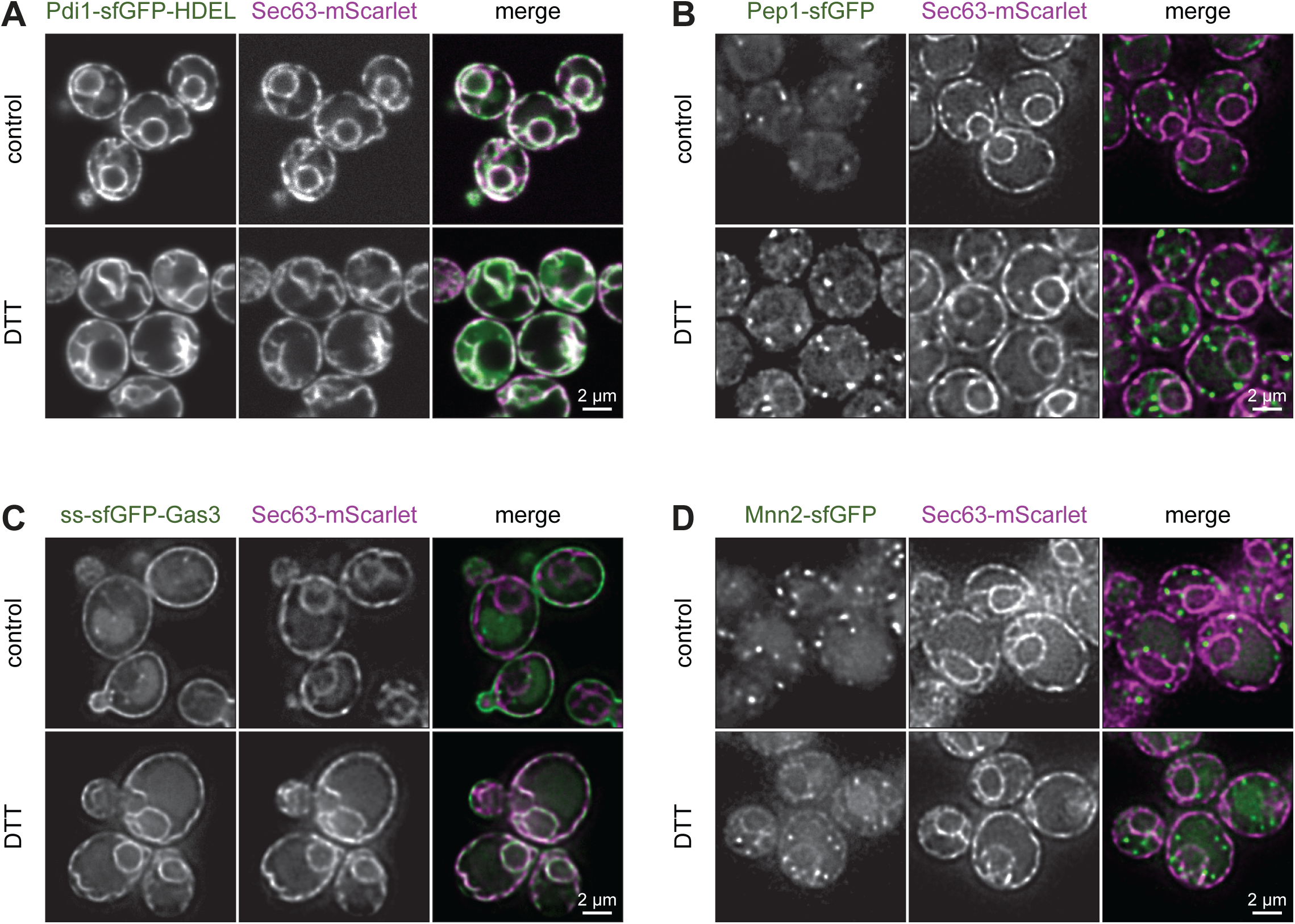
ER reflux and redistribution of vacuole, plasma membrane and Golgi proteins towards the ER upon ER stress. **(A)** Confocal fluorescence images of mid sections of control and DTT-treated cells expressing the ER marker Sec63-mScarlet and Pdi1-sfGFP-HDEL. **(B)** Deconvolved widefield fluorescence images of mid sections of control and DTT-treated cells expressing the ER marker Sec63-mScarlet and Pep1-sfGFP. **(C)** As in panel B but for ss-sfGFP-Gas3 (ss = signal sequence for ER entry). **(D)** As in panel B but for Mnn2-sfGFP. Pdi1 accumulates in the cytosol; Pep1, Gas3 and Mnn2 redistribute towards the ER.

**Figure S7.**
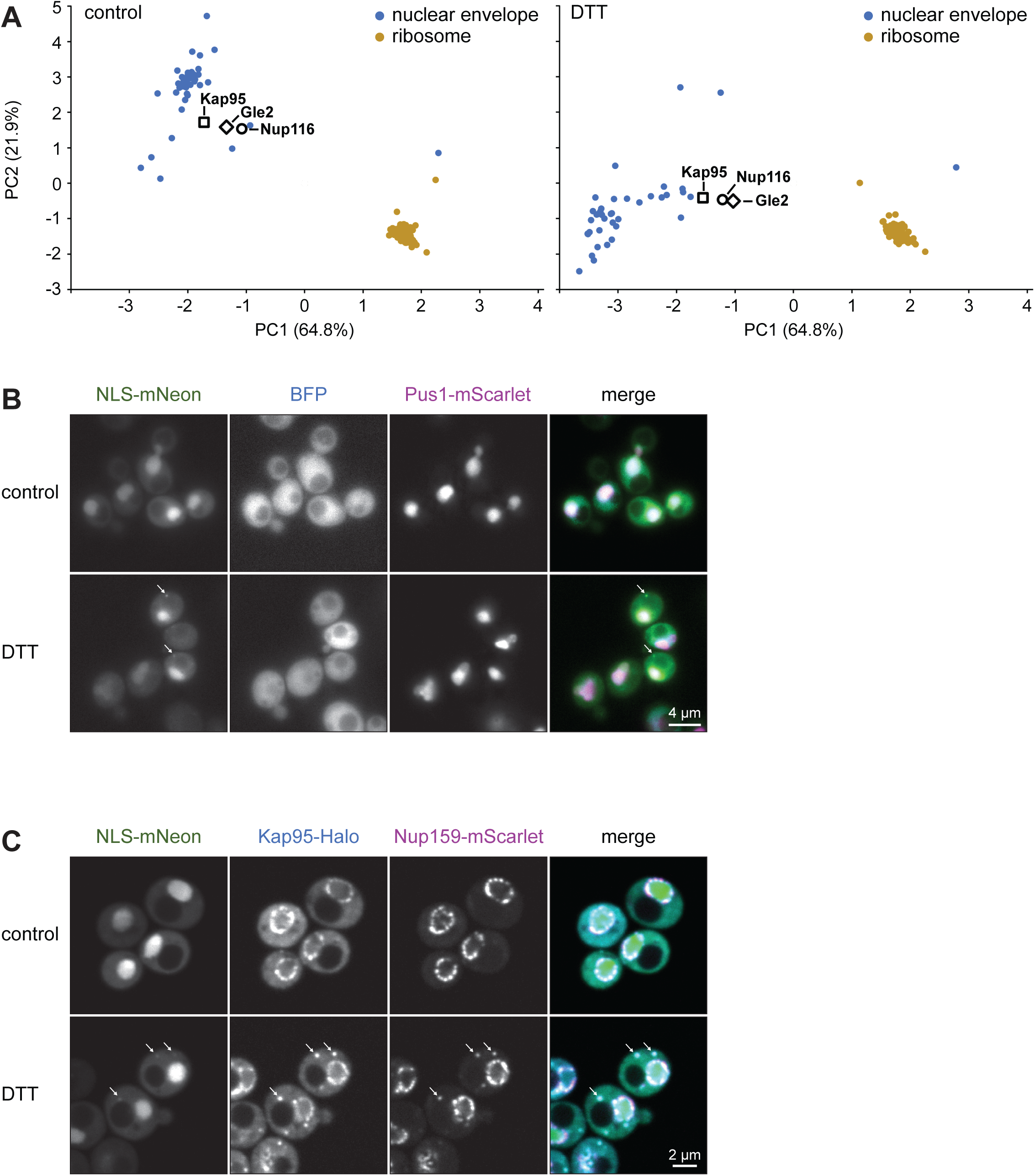
Redistribution of nucleoporins and importins and disturbed nuclear import upon ER stress. **(A)** PCA plot of organellar maps of control and DTT-treated cells highlighting the shifts of the nucleoporins Nup116 and Gle2 and the importins Kap95. The ribosome cluster is shown for reference. **(B)** Confocal fluorescence images of mid sections of control and DTT-treated cells. Cells expressed cytosolic BFP and the nuclear marker Pus1-mScarlet and additonally expressed mNeon fused to a nulcear localization sequence (NLS-mNeon) under the control of an estradiol-inducible promoter system. NLS-mNeon expression was induced for 50 min before imaging. Arrows indicate cytosolic NLS-mNeon puncta in DTT-treated cells. **(C)** As in panel B except that cells constitutively expressed Nup159-Scarlet and Kap95-Halo, and inducibly expressed NLS-mNeon.

