## Supplemental Dataset 3 for "Dynamic Organellar Mapping in yeast reveals extensive protein localization changes during ER stress": Notes for Supplemental Dataset 3.docx

This file contains all data required to recapitulate the DOM-ABC filtering, transformation and analysis of proteomic data generated in this study.

The data are split into two pools – the six steady-state maps (from untreated yeast), and the 3 x 3 ER stress maps (from control, DTT- and tunicamycin-treated cells).

Below is a step-by-step guide to upload and analyse the data.

**1. Steady-state maps**

Go to

<https://domabc.bornerlab.org/QCtool>

Click on:

**Format and analyse single experiment**

From the Supplemental Dataset 3 Folder, open the Steady State Maps folder. Select the text file

**Protein groups steady state maps**

Scroll down to ‘Load Settings’, and select

**domqc settings steady state maps**

Wait while the page updates itself, box by box.

In the box organelle markers, upload the custom file

**Yeast markers DOMABC_1908**

In the box annotation file, upload the custom file

**YeastGenes_stream**

(This step is not absolutely required, but generates the yeast gene names we originally used)

Press

**Run processing**

Wait for the analysis to finish.

Scroll down and you can now access the various data tabs for PCA, depth assessment, and download.

For PCA, de-select ‘fix aspect ratio by variability’, and select PC1 vs PC2.

In the download section, you can download the profiling data, or the complete analysis as a .json file.

To perform SVM analysis, scroll up to the top and select the

**Benchmark**

tab. The steady state maps dataset (called **YSM6**, for **yeast steady state maps with six replicates**) will already be loaded, so press

**Align and analyse selected datasets**

Wait for the analysis to finish.

Scroll down, and select the

**SVM analysis**

tab.

Select the

**Run SVMs**

tab.

Set test set proportion to 0.

Press

**Run training**

Be patient, it will take a while.

Once the training has finished, press

**Run predictions**

When done, select

**Download SVM predictions.**

Please note: If you downloaded the .json file from the ‘Format and analyse Single experiment’ section, you can also load this directly into the Benchmarking tool in future.

**2. Analysing the ER stress maps**

Go to

<https://domabc.bornerlab.org/QCtool>

Click on:

**Format and analyse single experiment**

From the Supplemental Dataset 3 folder, open the folder ‘ER Stress Maps’. Select the text file

**Protein groups ER stress maps**

Scroll down to ‘Load Settings’, and select

**domqc settings ER stress maps**

Wait while the page updates itself, box by box.

In the box organelle markers, upload the custom file

**Yeast markers DOMABC_1908**

In the box annotation file, upload the custom file

**YeastGenes_stream**

(This step is not absolutely required, but generates the yeast gene names we originally used)

Press

**Run processing**

Wait for the analysis to finish.

Scroll down and you can now access the various data tabs for PCA, depth assessment, and download.

You can download the profiling data, or the complete analysis as a .json file.

To perform the MR movement analysis, select the tab

**Movement analysis.**

To compare DTT treatment and control maps, just press

**Calculate/display MR plot**

This will take a while.

Scroll down to see the results.

To compare Tunicamycin treatment and control maps, select the corresponding datasets in the dialogue box near the top of the MR analysis tool, and run the analysis as above.

To perform SVMs and cross-dataset benchmarking, you need to repeat the steps above three times, but **for the individual control, DTT and tunicamycin protein groups datasets** provided here. So upload

**Protein groups ER stress maps CON**

with

**domqc settings ER stress maps CON**

and format as above;

upload

**Protein groups ER stress maps DTT**

with

**domqc settings ER stress maps DTT**

and format as above;

upload

**protein groups ER stress maps TUN**

with

**domqc settings for ER stress maps TUN**

and format as above.

Then go to the

**Benchmark**

Tab.

All three individual datasets will already have been loaded. Make sure that only these three datasets are selected, before pressing

**Align and analyse selected datasets**

Once the analysis is finished, you can access the SVM tool and other functionalities.

**Generating 7-datapoint profiles including the cytosolic fraction**

The data upload is as described above – ‘Format and analyse single experiment’ – the protein groups files (‘Steady state maps’ or ‘ER stress maps’) also include the cytosol data. There is only one extra step during set up. Before you press the ‘Run analysis’ button, go to the ‘Fractions’ button. In the empty box next to the last fraction, type ‘Cyt’. This will include the cytosolic fraction in the analysis. Now press ‘Run analysis’. Fraction weighting with protein yields and renormalization are currently performed outside DOM-ABC (see Methods for details).
