## Supplementary material for "Dynamic Organellar Mapping in yeast reveals extensive protein localization changes during ER stress": Table S7

**Table S7.** Plasmids used in this study. GEM, GAL4DBD-EstR-Msn2TAD; NLS, nuclear localization sequence; Kar2ss, Kar2 signal sequence.

| Plasmid | Alias | Source |
| --- | --- | --- |
| pFA6a-GFP(S65T)-HIS3 | pSS039 | Longtine, 1998 |
| pFA6a-mNeonGreen-klTRP1 | pSS904 | this study |
| pFA6a-mScarlet-I3-HIS3 | pSS1421 | this study |
| pFA6a-mScarlet-I3-klTRP1 | pSS1426 | this study |
| pFA6a-yeGFP-HDEL-kanMX6 | pSS451 | Young, 2013 |
| pFA6a-sfGFP-klTRP1 | pSS1499 | this study |
| pFA6a-mCherry-kanMX6 | pSS061 | this study |
| pFA6a-nat-P_CYC_-yeGFP | pSS1066 | Janke, 2004 |
| pFA6a-mNeonGreen-HIS3 | pSS447 | Papagiannidis, 2021 |
| pFA6a-HaloTag-klTRP1 | pSS1411 | this study |
| pRS415-P_TEF_ | pSS023 | Mumberg, 1995 |
| pRS406-P_GPD_-mCherry-Ubc6 | pSS117 | Benoit Kornmann |
| pRS406-P_TEF_-mCherry-Ubc6 | pSS482 | this study |
| pRS303H-P_GPD_-TagBFP | pSS349 | Szoradi, 2018 |
| pRS406-P_TEF_-TagBFP-Ubc6 | pSS988 | this study |
| pNH605-P_ADH_-GEM | pDEP151 | David Pincus |
| pFA6a-hph | pSS033 | Janke, 2004 |
| pFA6a-nat-P_GPD_-yeGFP | pSS440 | Janke, 2004 |
| pFA6a-hph-P_GPD_-yeGFP | pSS456 | this study |
| pRS405-P_GAL_-Kar2ss-sfGFP-HDEL | pSS1223 | this study |
| pFA6a-hph-P_GAL_-Kar2ss-sfGFP | pSS1367 | this study |
| pNH605-P_ADH_-GEM-P_GAL_ | pSS474 | Schmidt, 2019 |
| pNH605- P_ADH_-GEM-P_GAL_-NLS-mNeonGreen-T_CYC_ | pSS1563 | this study |
