## Supplementary material for "Dynamic Organellar Mapping in yeast reveals extensive protein localization changes during ER stress": Table S8

**Table S8.** Yeast strains used in this study. BFP, TagBFP; Cherry, mCherry; GEM, GAL4DBD-EstR-Msn2TAD; Kar2ss, Kar2 signal sequence; Neon, mNeonGreen; Scarlet, mScarlet-I3; sfGFP, superfolder GFP; yeGFP, yeast-enhanced GFP.

| Strain | Relevant genotype | Source |
| --- | --- | --- |
| SSY122 | *ADE2 leu2-3,112 trp1-1 ura3-1 his3-11,15 MATa* | Szoradi, 2018 |
| SSY4217 | *Sec63-Scarlet::HIS3* | this study |
| SSY4404 | *Sec63-Scarlet::HIS3 Pdi1-sfGFP-HDEL::kan* | this study |
| SSY4403 | *Sec63-Scarlet::HIS3 Sil1-sfGFP-HDEL::kan* | this study |
| SSY4356 | *Sec63-Scarlet::HIS3 Ero1-sfGFP::TRP1* | this study |
| SSY4355 | *Rtn1-Cherry::kan P_CYC_-yeGFP-Dpm1::nat Yop1-HaloTag::klTRP1 ura3::P_TEF_-BFP-Ubc6-URA3* | this study |
| SSY4220 | *Sec63-Scarlet::HIS3 Prc1-sfGFP::TRP1* | this study |
| SSY4279 | *Sec63-Scarlet::HIS3 Atg42-sfGFP::TRP1* | this study |
| SSY4278 | *Sec63-Scarlet::HIS3 Pep1-sfGFP::TRP1* | this study |
| SSY4251 | *Sec63-Scarlet::HIS3 leu2::P_ADH_-GEM-LEU2* | this study |
| SSY4280 | *Sec63-Scarlet::HIS3 leu2::P_ADH_-GEM-LEU2*  *P_GAL_-Kar2ss-sfGFP-Gas3::hph* | this study |
| SSY4281 | *Sec63-Scarlet::HIS3 leu2::P_ADH_-GEM-LEU2*  *P_GAL_-Kar2ss-sfGFP-Gas1::hph* | this study |
| SSY4283 | *Sec63-Scarlet::HIS3 leu2::P_ADH_-GEM-LEU2*  *P_GAL_-Kar2ss-sfGFP-Utr2::hph* | this study |
| SSY4259 | *Sec63-Scarlet::HIS3 Mnn2-sfGFP::TRP1* | this study |
| SSY4260 | *Sec63-Scarlet::HIS3 Mnn5-sfGFP::TRP1* | this study |
| SSY4394 | *Sec63-Scarlet::HIS3 Aur1-sfGFP::TRP1* | this study |
| SSY1212 | *ura3::P_GPD_-Cherry-Ubc6-URA3* | this study |
| SSY3788 | *ura3::P_GPD_-Cherry-Ubc6-URA3 Pom152-Neon::HIS3* | this study |
| SSY3789 | *ura3::P_GPD_-Cherry-Ubc6-URA3 Nup170-Neon::HIS3* | this study |
| SSY3790 | *ura3::P_GPD_-Cherry-Ubc6-URA3 Nup159-Neon::HIS3* | this study |
| SSY3791 | *ura3::P_GPD_-Cherry-Ubc6-URA3 Nup133-Neon::HIS3* | this study |
| SSY3792 | *ura3::P_GPD_-Cherry-Ubc6-URA3 Nup57-Neon::HIS3* | this study |
| SSY3793 | *ura3::P_GPD_-Cherry-Ubc6-URA3 Nic96-Neon::HIS3* | this study |
| SSY3794 | *ura3::P_GPD_-Cherry-Ubc6-URA3 Nup1-Neon::HIS3* | this study |
| SSY3885 | *ura3::P_GPD_-Cherry-Ubc6-URA3 Nup82-Neon::HIS3* | this study |
| SSY3886 | *ura3::P_GPD_-Cherry-Ubc6-URA3 Nup116-Neon::HIS3* | this study |
| SSY3980 | *ura3::P_GPD_-Cherry-Ubc6-URA3 Nup49-Neon::HIS3* | this study |
| SSY3981 | *ura3::P_GPD_-Cherry-Ubc6-URA3 Nsp1-Neon::HIS3* | this study |
| SSY4071 | *ura3::P_GPD_-Cherry-Ubc6-URA3 Nup42-Neon::HIS3* | this study |
| SSY4072 | *ura3::P_GPD_-Cherry-Ubc6-URA3 Gle1-Neon::HIS3* | this study |
| SSY4073 | *ura3::P_GPD_-Cherry-Ubc6-URA3 Gle2-Neon::HIS3* | this study |
| SSY4074 | *ura3::P_GPD_-Cherry-Ubc6-URA3 Dyn2-Neon::HIS3* | this study |
| SSY4129 | *ura3::P_GPD_-Cherry-Ubc6-URA3 Nup100-Neon::HIS3* | this study |
| SSY4410 | *ura3::P_GPD_-Cherry-Ubc6-URA3 Nup157-Neon::HIS3* | this study |
| SSY4411 | *ura3::P_GPD_-Cherry-Ubc6-URA3 Nup59-Neon::HIS3* | this study |
| SSY4413 | *ura3::P_GPD_-Cherry-Ubc6-URA3 Ndc1-Neon::HIS3* | this study |
| SSY4414 | *ura3::P_GPD_-Cherry-Ubc6-URA3 Nup53-Neon::HIS3* | this study |
| SSY4415 | *ura3::P_GPD_-Cherry-Ubc6-URA3 Pom34-Neon::HIS3* | this study |
| SSY4418 | *ura3::P_GPD_-Cherry-Ubc6-URA3 Nup188-Neon::HIS3* | this study |
| SSY4419 | *ura3::P_GPD_-Cherry-Ubc6-URA3 Nup84-Neon::HIS3* | this study |
| SSY4420 | *ura3::P_GPD_-Cherry-Ubc6-URA3 Nup120-Neon::HIS3* | this study |
| SSY4421 | *ura3::P_GPD_-Cherry-Ubc6-URA3 Mlp2-Neon::HIS3* | this study |
| SSY4446 | *ura3::P_GPD_-Cherry-Ubc6-URA3 Nup192-Neon::HIS3* | this study |
| SSY4456 | *ura3::P_GPD_-Cherry-Ubc6-URA3 Nup85-Neon::HIS3* | this study |
| SSY4457 | *ura3::P_GPD_-Cherry-Ubc6-URA3 Nup2-Neon::HIS3* | this study |
| SSY4458 | *ura3::P_GPD_-Cherry-Ubc6-URA3 Nup60-Neon::HIS3* | this study |
| SSY4459 | *ura3::P_GPD_-Cherry-Ubc6-URA3 Mlp1-Neon::HIS3* | this study |
| SSY4463 | *ura3::P_GPD_-Cherry-Ubc6-URA3 Nup145C-Neon::HIS3* | this study |
| SSY4293 | *ura3::P_GPD_-Cherry-Ubc6-URA3 Kap95-Neon::HIS3* | this study |
| SSY4444 | *ura3::P_GPD_-Cherry-Ubc6-URA3 Nup116-Neon::HIS3 Kap95-HaloTag::klTRP1* | this study |
| SSY4571 | *trp1::P_TEF_-BFP-Ubc6-TRP1 Nup159-Scarlet::HIS3 Kap95-HaloTag::kan leu2::P_ADH_-GEM-P_GAL_-NLS-Neon-T_CYC_-LEU2* | this study |
| SSY4574 | *his3::P_GPD_-BFP-hph leu2::P_ADH_-GEM-P_GAL_-NLS-Neon-T_CYC_-LEU2 Pus1-Scarlet::klTRP1* | this study |
